# AMPK acts to remove immune barriers to CD8^+^ T cell-mediated immunity against hepatocellular carcinoma

**DOI:** 10.1101/2024.12.16.628539

**Authors:** Hui-Hui Hu, Xuefeng Wang, Bin Lan, Haili Cheng, Hong Wen, Fangfang Chen, Jianfeng Wu, Mengqi Li, Jiazhou Chen, Dongxu Chen, Shiyu Lin, Mingyang Yang, Zhenhua Wu, Zhong-Zheng Zheng, Fuqing Chen, Jianyin Zhou, Gang Chen, Yu Chen, Xianming Deng, Chen-Song Zhang, Jingfeng Liu, Sheng-Cai Lin

## Abstract

Dysregulated metabolism in tumor tissues, and para-tumor tissues alike, can lead to immunosuppression, which may underlie cancer development. However, metabolic intervention as a therapeutic strategy has been of no avail. In this study, we explored the anti-cancer therapeutic effect of aldometanib that specifically targets lysosome-associated aldolase to mimic glucose starvation and thereby activates the lysosomal AMP-activated protein kinase (AMPK), a master regulator of metabolic homeostasis. We show that aldometanib inhibits the growth of HCC in an AMPK-dependent manner, allowing hepatoma-bearing mice to survive to mature ages, although aldometanib does not possess cytotoxicity towards HCC or normal cells. Intriguingly, aldometanib exerts anti-cancer effects only in immune-competent host mice, but not in immune-defective mice. We have further found that the HCC tissues in aldometanib-treated mice are massively infiltrated with CD8^+^ T cells, which is not seen in mice with liver-specific knockout of *AMPKα*. Our findings thus suggest that the metabolic regulator AMPK rebalances the tumor microenvironment to allow the cytotoxic immune cells inside the body to eliminate cancer cells and effectively contain the tumor tissues. The finding that metabolic intervention can make cancer a life-long manageable disease, may potentially usher in a new era of cancer therapy.

---

Emerging evidence suggests that metabolic imbalance contributes to the development of cancers (*1, 2*). Dysregulated metabolism leads to altered compositions of metabolites and properties of proteins inside the cellular compartments, as well as in the extracellular tumor microenvironment (TME) between the tumor and para-tumor tissues (*3*). One of the cancers that have been extensively studied for association with metabolic alterations is hepatocellular carcinoma (HCC) (*4, 5*). Metabolic alterations in glycolysis, the TCA cycle, nucleotide synthesis, amino acid metabolism, de novo lipogenesis, and cholesterol synthesis in HCC cells, have all been implicated in the formation and development of HCC (*4*). Accumulation of lactate, for instance, may acidify the TME, and has been shown to contribute to the establishment of an immunosuppressive microenvironment of para-HCC tissues by reducing levels of chemokines such as CXCL9 and CXCL10, which are known to facilitate the infiltration of the tumouricidal CD8^+^ T cells (*6*). Another possible mechanism by which lactate exerts pro-tumorigenic effects is through lysine lactylation of various metabolic enzymes (*7*). Tremendous efforts have hence been made to identify chemical compounds that target various metabolic routes in search of effective therapies to treat cancers including HCC, and some of the identified compounds have been undergoing clinical trials (*4, 5*). For the same reason, AMPK, which is a master regulator of metabolic homeostasis controlling the overall metabolic network (*8–10*), has also been targeted for screening clinically useful drugs for treating HCC (*11, 12*).

AMPK is activated in response to physiological declines of glucose (*13*), decrease of energy levels reflected by increased ratios of AMP:ATP or ADP:ATP (*14*), or global changes of cellular Ca^2+^ levels (*15*). Upon activation, AMPK phosphorylates a series of substrates to slow down anabolic activities such as the synthesis of fatty acids, cholesterol, and proteins, which are critical constituents of the building blocks for the proliferation of normal cells and cancer cells as well (*11, 16*). It has been shown that statin-mediated inhibition of HMG-CoA reductase, a classic substrate of AMPK, to lower the synthesis of cholesterol, can attenuate the development of HCC in patients (*17–19*). Similarly, the compound ND-654 inhibits the development of HCC by inhibiting acetyl-CoA carboxylases (ACC, for de novo synthesis of fatty acid), another classic substrate of AMPK (*20*). Not surprisingly, endeavors have been made to develop pharmacological agonists directly targeting AMPK for potential efficacy in treating HCC. However, due to the indiscriminate activation of all subcellular pools of AMPK by such identified chemicals, most of the compounds may have adverse effects (*21*). For example, the pan-AMPK activator MK-8722 causes cardiac hypertrophy, although it robustly activates AMPK and lowers blood glucose (*22*). Similarly, although the mitochondrial inhibitors activate AMPK via elevating AMP:ATP or ADP:ATP ratios, the inhibition of mitochondria per se may be harmful, particularly under prolonged treatments (*23, 24*). Nevertheless, it is encouraging to see that the AMPKβ1-specific agonists with acceptable safety profiles, have shown effects in alleviating liver diseases such as fatty liver and nonalcoholic steatohepatitis (NASH) (*25–27*).

We have previously delineated a pathway that senses declines of glucose, seen during situations that occur during physiological situations such as fasting, and transmits the low-glucose signal to the activation of AMPK on the surface of the lysosome (*28–31*), and importantly, the concomitant inhibition of the pro-anabolic mTORC1 (*29, 32*). This glucose sensing-AMPK pathway involves a direct sensing of the availability of fructose-1,6-bisphosphate (FBP), a glycolytic intermediary metabolite that is a substrate of the glycolytic aldolase (*30*). Aldolase, when not occupied by FBP, acts to inhibit the endoplasmic reticulum-localized cation channels TRPV, which themselves, after the inhibition, physically interact with and inhibit the lysosomal proton pump v-ATPase (*30, 31*). Inactivated v-ATPase, along with conformational changes of its associated proteins, including the Ragulator complex, solicits translocation of the AXIN:LKB1 complex to the surface of the lysosome, whereby the lysosomally localized AMPK is phosphorylated by the co-translocated LKB1 and becomes activated (*29*). This route for activating the lysosomal pool of AMPK also offers a spatiotemporal regulation of metabolism (*33*), offering targets for identifying compounds that specifically activate the lysosomal AMPK to mimic fasting states and elicit beneficial effects. The benefits of AMPK activation via this lysosomal pathway can be reflected by the fact that metformin at clinical doses takes a ride on this pathway to elicit various effects, including alleviation of fatty liver (*34–36*). In fact, metformin has been shown to alleviate the risk of HCC development and progression in some clinical studies ((*37–40*); phase III clinical trial NCT03184493).

By using aldolase as a target, we have identified an inhibitor termed aldometanib, which specifically prevents the v-ATPase-associated aldolase, but not the cytosolic aldolase engaged in glycolysis, from binding to FBP even in high glucose, thereby mimicking a state of low glucose to activate AMPK via the lysosomal pathway (*41*). In an AMPK-dependent manner, aldometanib alleviates hyperglycemia, fatty liver, and NASH in obese rodents (*41*).

In this study, we wondered whether aldometanib could possibly treat HCC, by correcting the metabolic dysregulation that might have been aggravated by pathological situations such as hepatitis or fatty liver, most prevalent pre-conditions for developing liver cancers. Notwithstanding that aldometanib does not exert cytotoxicity towards cells, we show that aldometanib drastically inhibits the growth and development of HCC in the DEN-treated and HFD (high-fat diet)-fed mice, in mice with hepatic knockout of *Trp53* coupled with overexpression of Myc (*MYC*;*Trp53*^−/−^ HCC (*42*)), and in xenografts established by HCC cell lines. Remarkably, treatment with aldometanib permits the HCC-bearing DEN-HFD mice to live to mature ages. We further found that treatment of aldometanib induces CD8^+^ cytotoxic T cells to massively surround and infiltrate into the tumor tissues, but not in the liver with knockout of AMPK. These observations suggest that aldometanib activates AMPK to revert the tumor environments to a situation that allows the migration of immune cells, thereby acting as a leverage for the immune system of the body to defend against cancer development to allow cancer-bearing mice to live to maturity.

## Aldometanib treatment allows HCC-bearing mice to live to normal ages

Mice were treated with diethylnitrosamine (DEN) at 4 weeks old, and fed with a HFD starting at week 7 of age (the DEN-HFD HCC (*43*); depicted in fig. S1A). From 40 weeks old (35 weeks after DEN injection), the DEN-HFD mice were treated with aldometanib dissolved in drinking water. We first titrated for effective concentrations of aldometanib on AMPK activation in the liver tissues, both HCC and para-HCC tissues, of these mice, and found that concentrations ranging from 50 to 150 mg/L effectively activated AMPK (as determined by the levels of the phosphorylation of Thr172 of AMPKα and phosphorylation of Ser79 of ACC1; fig. S1A), and concomitantly inhibited mTORC1 (determined by the phosphorylation of the mTORC1 substrate S6K at its Thr389 site; fig. S1A). We also found that aldometanib at such doses did not affect the energy levels, as evidenced by the unchanged ratios of AMP:ATP or ADP:ATP in both HCC and adjacent noncancerous (fig. S1B), consistent with the previous report that aldometanib acts to activate AMPK via the lysosomal pathway, independently of the increase of AMP (*41*).

We then determined the effects of aldometanib on the lifespan of the DEN-HFD mice. Strikingly, aldometanib at 100 mg/L extended the survival time of the HCC-bearing mice to normal ages compared to the control group (Fig. 1A and table S1), with the treated HCC mice having a median lifespan of approximately 805 days, which is even slightly longer than the median surviving time (796 days) of the healthy, untreated mice (Fig. 1A; and ref. (*41, 44*)). Aldometanib also reduced the sizes and weights of the HCC tumors (Fig. 1B and fig. S1, C to F; collected at 48 weeks of age as a representative time point). The suppression of HCC by aldometanib displayed a dose dependence, with mild inhibition at 50 mg/L to strong at 150 mg/L (Fig. 1B and fig. S1, C to F). Importantly, aldometanib administered at late stages, as late as 30 weeks after DEN injection, by the time tumors had already formed (fig. S2A; see also ref. (*43*)), still resulted in strong inhibition of HCC (Fig. 1C and fig. S2, B to F). Reflecting improved liver damage, alanine aminotransferase (ALT) and aspartate transferase (AST) in the serum of DEN-HFD mice treated with aldometanib (fig. S1G and S2G) were significantly reduced. In addition, the staining of alpha fetal protein (AFP), an indicator of the severity and metastasis of liver cancer (*45*), in the HCC tumor sections, was also decreased (Fig. 1D). We also found a significant reduction of lipid droplets and triglyceride in the liver of aldometanib-treated DEN-HFD mice (fig. S3, A and B), consistent with AMPK being activated by the drug. These results indicate that aldometanib not only effectively contains HCC, but also permits the HCC mice to live to mature ages.

**Fig. 1.**
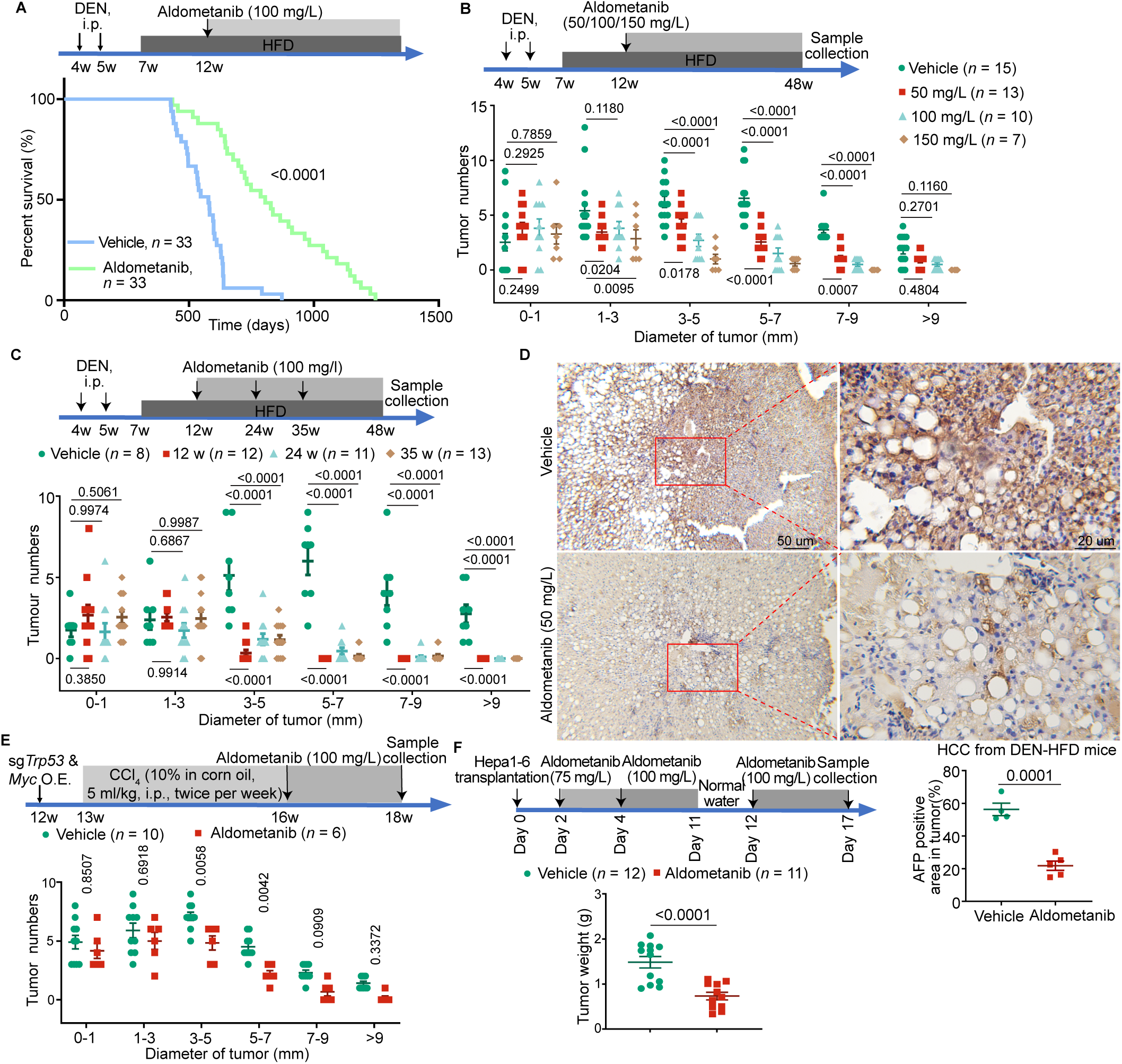
Aldometanib inhibits hepatocellular carcinomas. **(A)** Aldometanib treatment allows HCC-bearing mice to live to a similar age to normal mice. Wildtype C57BL/6 mice (4 weeks old) were intraperitoneally injected with DEN once a week for 2 weeks, followed by feeding with HFD 2 weeks later. The mice were then administered with aldometanib at 100 mg/L in drinking water starting at 12 weeks (see also statistical analyses in table S1). The average survival age of the aldometanib-treated DEN-HFD mice is 805 days, similar to 795 days for normal healthy mice as determined by us (*41*) and others (*44*). **(B)** Aldometanib inhibits the growth of HCC in DEN-HFD mice. Mice were induced to develop HCC using DEN and HFD as in (A). Aldometanib at 50, 100, or 150 mg/L was administered in drinking water starting at week 12 of age, and the tissue samples were collected at week 48 (depicted in the upper panel). Statistics of the numbers of tumors in each size/diameter: 0–1 mm, 1–3 mm, 3–5 mm, 5–7 mm, 7–9 mm, or > 9 mm (lower panel; shown as means ± s.e.m., *n* values indicate the number of mice used for each condition and are labeled in each panel; and *P* values were calculated by two-way ANOVA, followed by Tukey’s test) for each dosage of aldometanib are shown. **(C)** Aldometanib effectively inhibits late-stage HCC. Mice were induced to develop HCC using DEN and HFD as in (A). Aldometanib at 100 mg/L was administered in drinking water starting at 12, 24, or 35 weeks old (depicted in the upper panel). Statistics of the numbers of tumors in each size/diameter (lower panel; shown as means ± s.e.m., *n* represents the number of mice for each starting age, and are labeled in each panel, with *P* values calculated by two-way ANOVA, followed by Tukey’s test) for each entry time of aldometanib are shown. **(D)** Aldometanib suppresses alpha fetal protein (AFP) levels in DEN-HFD mice. HCC tissues from both the aldometanib-treated and untreated DEN-HFD mice, prepared as in (B), were subjected to immunohistochemistry staining for AFP. Representative images at different magnifications are shown on the upper panel, and the percentages of AFP-positive area in the HCC tissues were calculated and are shown on the lower panel (means ± s.e.m., *n* = 4 (vehicle) or 5 (aldometanib) mice for each treatment, with *P* values calculated by two-sided Student’s *t*-test). **(E)** Aldometanib inhibits HCC in *MYC*;*Trp53*^−/−^ mice. Wildtype BALB/c mice (12 weeks old) were hydrodynamically injected, via tail vein, with plasmids carrying sgRNAs targeting *Trp53* for knockout, along with the plasmid for overexpressing Myc. One week later, mice were then intraperitoneally injected with CCl_4_ twice a week for 3 weeks (depicted in the upper panel). Aldometanib was administered at 100 mg/L in drinking water starting at 16 weeks old. At week 18, mice were euthanized, and the statistics of the numbers of tumors in each size/diameter (lower panel; shown as means ± s.e.m., n numbers are labeled in each panel, with *P* values calculated by two-way ANOVA, followed by Sidak’s test) are shown. **(F)** Aldometanib reduces the sizes of xenografts derived from the Hepa1-6 hepatoma cells. Hepa1-6 cells were transplanted into the left liver lobe of C57BL/6 mice to form xenografts. At day 2 post-transplantation, the mice were treated with aldometanib at 100 mg/L in drinking water for 15 days (depicted in the upper panel). At day 17, mice were euthanized, and the weights of the xenografts were determined (lower panel; shown as means ± s.e.m., n represents the number of mice, and are labeled in each panel, with *P* values calculated by two-sided Student’s *t*-test). Experiments in this figure were performed three times.

We also tested the efficacy of aldometanib on another HCC model in that *Trp53* is knocked out and Myc is overexpressed, two of the most frequently altered genes in HCC patients, in the mouse liver (*MYC*;*Trp53*^−/−^ HCC (*42*)). Unlike the DEN-HFD HCC, the *MYC*;*Trp53*^−/−^ HCC did not involve the process of fatty liver formation (*42*). As shown in Fig. 1E and fig. S4, A to F, aldometanib also suppressed the development of HCC in the *MYC*;*Trp53*^−/−^ mice. We also generated xenografts derived from the hepatoma cell line Hepa1-6 (*46*) transplanted in the left lobe of the mouse liver. We found that aldometanib also significantly reduced the sizes of the xenografts (Fig. 1F and fig. S5, A to F), especially given that the drug was applied after the formation of the xenograft tumors. The data above indicate that aldometanib showed strong effects on different HCC models including the reduction of the tumor sizes and wellness of the mice, with the survival of HCC mice to mature ages as the most important efficacy endpoint.

## AMPK in non-cancerous host liver plays a dominant role in suppression of HCC by aldometanib

We next investigated the role of AMPK in the aldometanib-mediated suppression of HCC in mice. First of all, we generated DEN-HFD mice with liver-specific double knockout of *AMPKα1* and *AMPKα2* (*AMPKα*-LKO, generated as in ref. (*35*)). The DEN-HFD treatment induced formation of HCC in the *AMPKα*-LKO mice at a similar onset to that in AMPK-wildtype mice (Fig. 2B and fig. S6A). Interestingly, aldometanib did not suppress the development of HCC in the liver lacking AMPK (Fig. 2, A and B, and fig. S6, A to E), indicating that aldometanib suppresses HCC progression indeed via activating AMPK. To differentiate the role of AMPK in the HCC tissue from that in the noncancerous para-tumor liver tissue, we examined the effect of aldometanib on the xenografts derived from *AMPKα*^-/-^ Hepa1-6 cells. We found that aldometanib could effectively inhibit the xenograft tumor of *AMPKα*^-/-^ Hepa1-6 cells (validated in fig. S6F) transplanted in the wildtype liver (Fig. 2, C and D and fig. S6, G to K), but not the xenografts grown in the host liver lacking AMPKα (*AMPKα*-LKO; Fig. 2, E and F, and fig. S6, L to P). In contrast, aldometanib only marginally impeded the growth of the xenografts of either wildtype Hepa1-6 cells when transplanted in the *AMPKα*-LKO host liver (Fig. 2, G and H, and fig. S6, Q to U). These results indicate that AMPK in the noncancerous para-tumor tissues plays a necessary and dominant role in the tumor suppression of aldometanib.

**Fig. 2.**
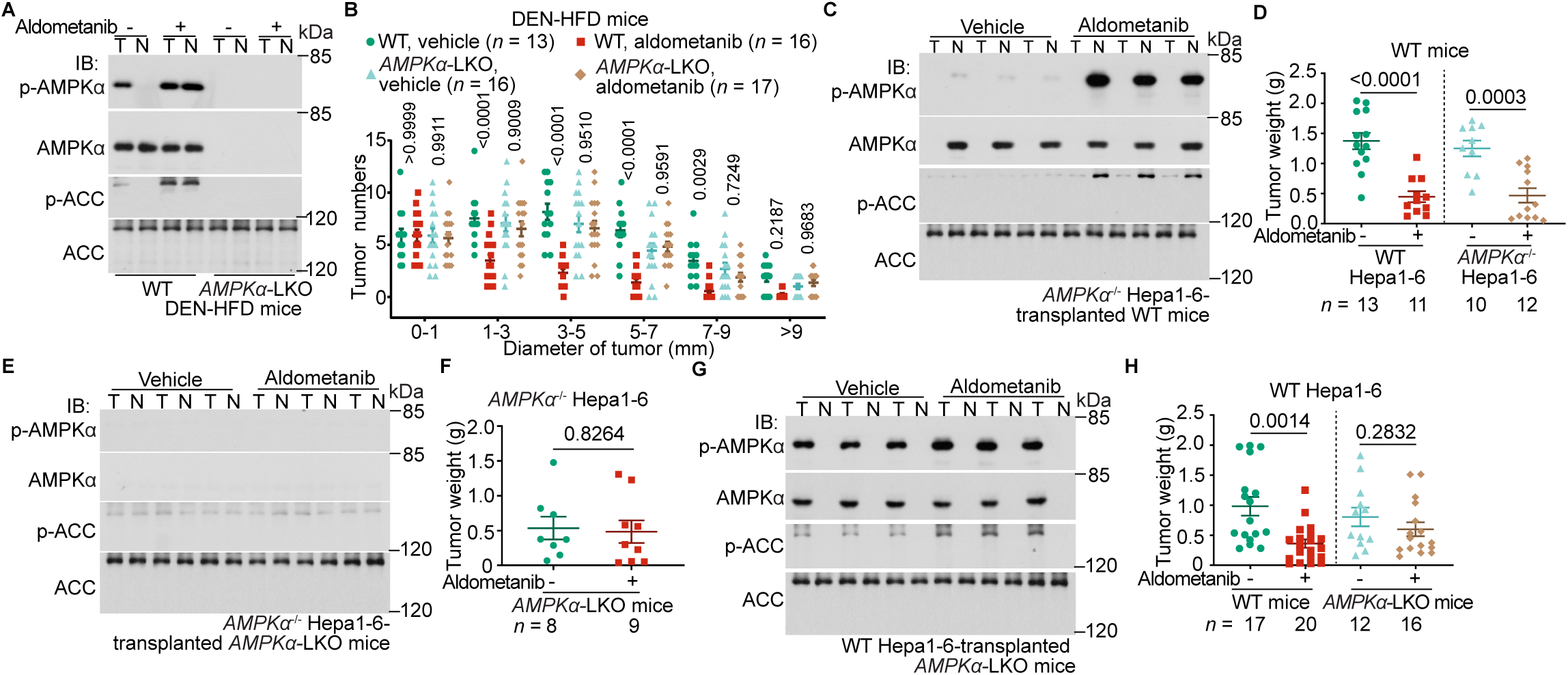
AMPK in para-HCC plays a dominant role in aldometanib-induced suppression of HCC. (**A** and **B**) Ablation of AMPK in the liver blocks the HCC-suppressing effects of aldometanib. The C57BL/6 mice with liver-specific AMPKα knockout (*AMPKα*-LKO) and wildtype mice were induced to develop HCC using DEN and HFD, followed by treatment with aldometanib as in Fig. 1A. AMPK activity in the liver (A) and the numbers of tumors in each size/diameter were determined ((B); shown as means ± s.e.m., *n* represents the number of mice, and are labeled in each panel; and *P* values calculated by two-way ANOVA, followed by Tukey). (**C** to **H**) Host liver AMPK plays a dominant role in the suppression of HCC xenografts by aldometanib. The wildtype ((C) and (D)) or *AMPKα*-LKO C57BL/6 mice ((E) and (F)) were transplanted with wildtype ((G) and (H)) or *AMPKα*^-/-^ ((C) to (F)) Hepa1-6 cells, followed by treatment with aldometanib as in Fig. 1F. AMPK activation ((C), (E), (G)) and tumor weights ((D), (F), (H); means ± s.e.m., *n* represents the number of mice, and are labeled in each panel; and *P* values calculated by two-sided Student’s *t*-test ((D), (F), and right panel of (H)) or by two-sided Student’s *t*-test with Welch’s correction (left panel of (H)) were then determined. Experiments in this figure were performed three times.

## Aldometanib does not possess cytotoxicity

We then explored how aldometanib inhibits the growth of HCC to allow the life-long survival of the HCC mice. We first examined whether aldometanib inhibits the growth of cells in vitro, by taking advantage that, unlike the mouse primary hepatocytes, BNL cells derived from normal liver tissue are able to proliferate in culture. We found that aldometanib showed inhibitory effects on the proliferation of BNL cells at a concentration of 50 nM (fig. S7C), which is similar to the concentrations detected in the liver of DEN-HFD HCC mice or in xenografts grown in normal mice (fig. S7, A and B) after administration of aldometanib at 100 mg/L. Aldometanib inhibited the growth of various other HCC cells, including Hepa1-6, JHH-7, and Huh7 (fig. S7C), with the growth inhibition of primary mouse HCC cells insignificant (fig. S7C). In line with the inhibition of the proliferation of these cells, the growth of HCC xenografts derived from Hepa1-6 cells were also inhibited (Fig. 3A). We also determined the cytotoxic activity of aldometanib to these cells in vitro, and surprisingly found that aldometanib did not cause the death of HCC cells, either apoptotic or necroptotic, even at 5,000 nM, a concentration much higher than the effective doses detected in the liver of aldometanib-treated mice (Fig. 3, B to E and fig. S7, D to H). Aldometanib also had no cytotoxicity towards normal hepatocytes (Fig. 3, F and G and fig. S7I). However, when HCC tissues of the aldometanib-treated mice were examined in vivo by TUNEL assays to detect cell death, a robust increase of the TUNEL signal was observed (Fig. 3H). Consistently, the levels of apoptotic markers, including cleavage of caspase-7 and the protein levels of PUMA, were increased in the aldometanib-treated HCC tissues (Fig. 3I). In addition, pyroptosis was also increased, as evidenced by the elevated levels of the cleaved form of GSDME (Fig. 3I). The lack of cytotoxicity in cultured cells notwithstanding, these data demonstrate that aldometanib is able to cause cell death inside the HCC grown inside the mouse, implying that the drug can induce cytotoxicity to tumor tissues by an innate mechanism.

**Fig. 3.**
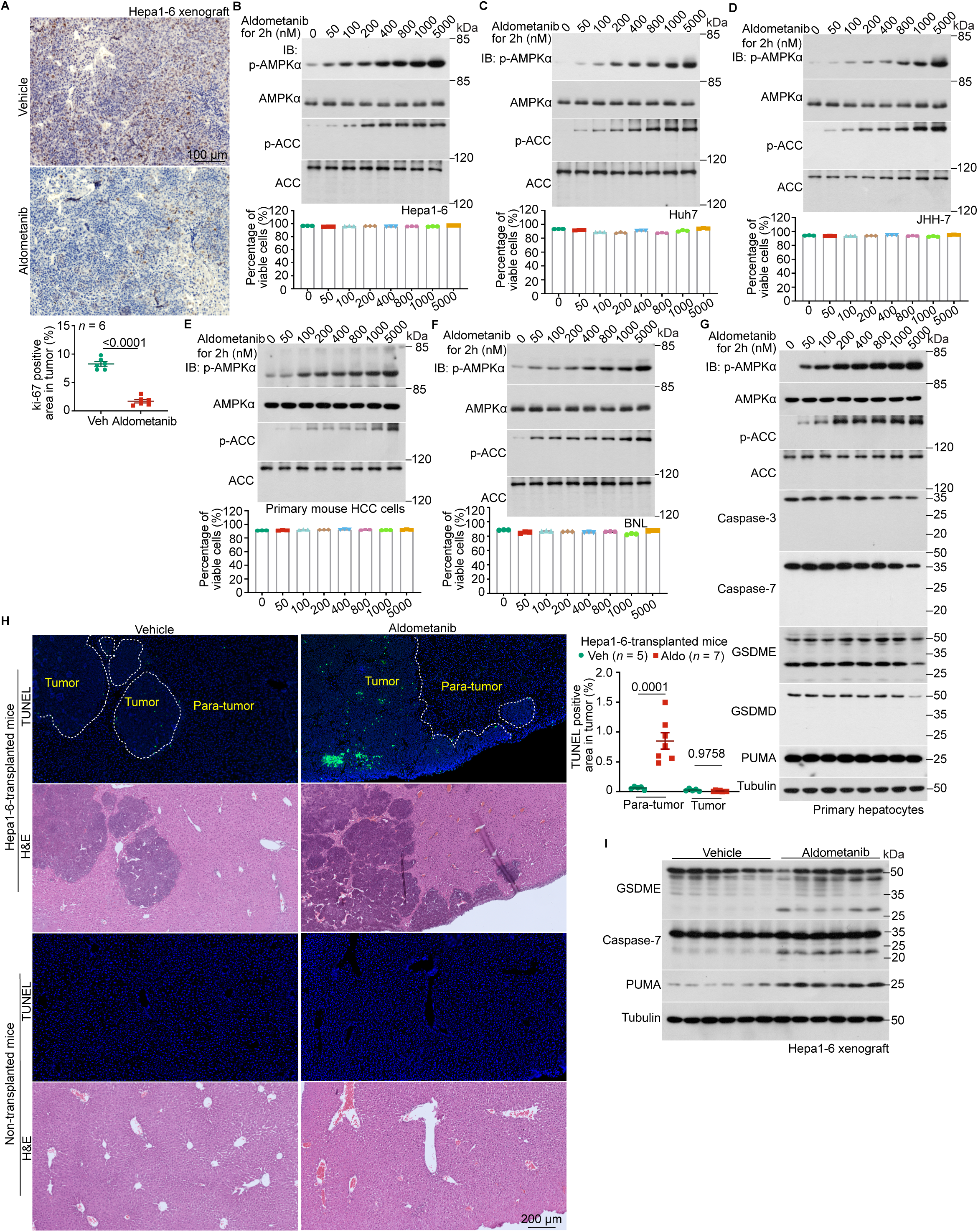
Aldometanib induces cytotoxicity specifically towards HCC tissues in vivo. (**A**) Aldometanib reduces the sizes of HCC xenografts derived from Hepa1-6 cells. HCC sections of the xenografts derived from Hepa1-6 cells, generated as in Fig. 1F, were subjected to immunohis-tochemistry staining for Ki-67. Representative images are shown on the left, and the percentages of Ki-67-positive areas within the HCC tissues were calculated and are shown on the right (means ± s.e.m., *n* = 6 mice for each treatment, with *P* values calculated by two-sided Student’s *t*-test). The scale bars are 100 μm. (**B** to **G**) Aldometanib does not exhibit cytotoxicity towards liver cells in culture. Various liver cell types, including the mouse HCC cell line Hepa1-6 (B), human HCC cell lines Huh7 (C) and JHH-7 (D), primary mouse HCC cells (E), as well as the normal mouse liver cell line BNL (F) and primary mouse hepatocytes (G), were treated with aldometanib at indicated concentrations for 2 h. The percentages of viable cells (calculated by subtracting early apoptotic, late apoptotic, and necroptotic cells from the total cell count) were determined using flow cytometry ((B) to (F); see gating strategy in fig. S7D, and representative density plots in fig. S7, D to I); data are shown at the bottom of each panel as means ± s.e.m., *n* = 3 biological replicates for each condition). See also immuno-blots of AMPK activation on the upper panels of (B) to (F), and the apoptotic and pyroptotic markers in (G). (**H** and **I**) Aldometanib induces cell death inside HCC tissues. HCC tissues from the xenograft derived from Hepa1-6 cells, generated as in Fig. 1F, were subjected to TUNEL assays ((H); representative images are shown on the left panel, and the percentages of TUNEL-positive area in the tumor or para-tumor tissues were calculated and are shown on the right panel as means ± s.e.m., *n* represents the number of mice, and are labeled in each panel, with *P* values calculated by two-way ANOVA, followed by Tukey), or immunoblotting for apoptotic and pyroptotic markers (I). The scale bars are 200 μm. Experiments in this figure were performed three times.

## Aldometanib induces infiltration of CD8^+^ T cells into tumors

We then determined how aldometanib-induced AMPK activation inhibits the development of the HCC tumors inside the mouse. As AMPK is a master regulator of metabolism, its activation may improve the composition of the constituents in the microenvironment between the noncancerous and tumor tissues, as well as in the extracellular fluids (*47*), which can generate an immune barrier. It is well-established that the cytotoxic lymphocytes, particularly the CD8^+^ T lymphocytes, act to induce the death of cancerous cells (*48*). To explore whether aldometanib affects the association with and infiltration of lymphocytes into the tumors, we next examined CD8^+^ T cells in the HCC tissues from the DEN-HFD mice or xenografts of Hepa1-6 cells, in wildtype or *AMPKα*-LKO host liver. In both of the HCC tissues, we found that aldometanib drastically enriched CD8^+^ T cells in the tumors grown in wildtype host liver (Fig. 4, A to D and fig. S8, A to D), but not in the *AMPKα*-LKO host liver (Fig. 4, C and D). In comparison, CD8^+^ T cells were rarely seen in the HCC from the vehicle (aldometanib-untreated) group (Fig. 4, A to D and fig. S8, A to D). In addition, knockout of *AMPKα* in the host liver blocked such induced enrichment of CD8^+^ T cells in the xenografts from either wildtype or *AMPKα*^-/-^ Hepa1-6 cells (Fig. 4D), while the depletion of *AMPKα* in the Hepa1-6 xenografts did not abolish the enrichment of CD8^+^ T cells (Fig. 4D). Increased levels of granzyme B and IFNγ in the CD8^+^ T cells were observed in the aldometanib-treated HCCs (Fig. 4, E to H and fig. S8, E to H), indicative of enhanced tumouricidal effects corroborative with the enrichment of the CD8^+^ T cells. Consistent with the increased presence of CD8^+^ T cells, we also observed a significant decrease in neutrophils and tumor-associated macrophages (TAMs) in the HCCs in mice treated with aldometanib (Fig. 4, I and J and fig. S8, I to K, O, and P), both of which are known to act as major immune barriers in the tumor microenvironment preventing the activation and infiltration of CD8^+^ T cells (*49, 50*). In addition, the numbers of lymphoid dendritic cells (DCs) and the expression levels of major histocompatibility complex II (MHC II) in myeloid DCs (*51, 52*), required for the activation (priming) of CD8^+^ T cells through the stimulation of the Th1 subtype of CD4^+^T cells (*53, 54*), were also increased with aldometanib treatment (Fig. 4, K and L, and fig. S8, L to N). We also observed that aldometanib increased to some extent the recruitment of Th2 cells, which are known to create an immunosuppressive environment in HCC (*55, 56*)—in 4 out of 18 samples analyzed (fig. S8Q). However, it is unlikely that the minor increase of Th2 cells would suppress CD8^+^ T cells, because the number of Th2 cells was approximately 30-fold lower than that of Th1 cells, which is insufficient to significantly alter the Th2:Th1 ratio in the HCC tissues (fig. S8Q). Furthermore, we found that aldometanib also promoted the infiltration of B cells (fig. S8R) and the γδT cells (fig. S8S), both of which are known to help the recruitment and activation of CD8^+^ T cells in HCC (*57–61*). Importantly, we found that when CD8^+^ T cells in mice were depleted by repeated intravenous injection of its neutralizing antibody, the aldometanib-enhanced enrichment of CD8^+^ T cells and suppression of HCC were significantly diminished (Fig. 4M and fig. S9, A to G). Furthermore, we observed that the suppression of HCC by aldometanib was greatly hindered in JHH-7 or Huh7 xenografts grown in the liver of immunocompromised BALB/c nude mice lacking CD8^+^ T cells (fig. S10, A to L). Similarly, when cells from human HCC tissues were transplanted into the liver of NOD-SCID mice, the tumor-suppressing effects of aldometanib were observed only in mice intraperitoneally injected with human peripheral blood mononuclear cells (PBMCs) containing CD8^+^ T cells, from healthy donors (Fig. 4N and fig. S10, M to R).

**Fig. 4.**
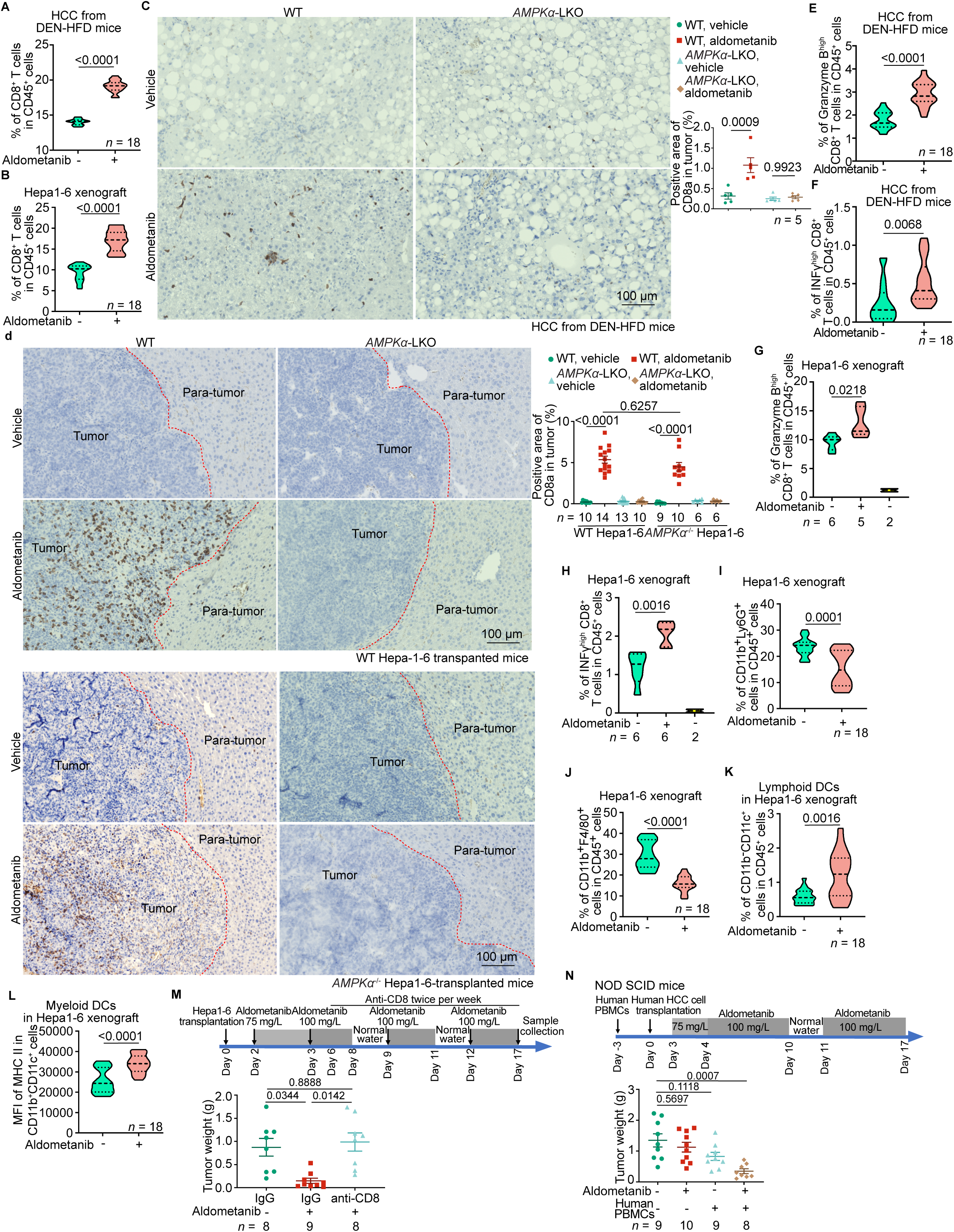
Aldometanib induces infiltration of CD8^+^ T cells into tumors. (**A** and **B**) Aldometanib promotes the infiltration of CD8^+^ T cells into HCC tissues. The HCC tissues from DEN-HFD mice ((A); prepared as in fig. S8A and collected at 40 weeks old) or Hepa1-6-derived xenografts ((B); generated as in fig. S8B and collected at day 10 post-transplantation), were digested with type I collagenase, followed by staining with CD8a-Alexa Fluor 488, CD4-APC, and CD45-PerCP-Cy5.5 (A) or CD8a-APC and CD45-Alexa Fluor 488 (B) antibodies to label the CD8^+^ T cells. The stained cells were then subjected to flow cytometry analysis for quantification of CD8^+^ T cells. Statistical data are shown as violin plots, in which the lower and upper dashed lines represent the first and the third quartile scores, respectively, the center dashed line represents the median, and the lower and upper limits denote minimum and maximum scores, respectively, and the same hereafter for all violin plots. *n* = 18 samples from 6 mice in each condition, with *P* values calculated by two-sided Student’s *t*-test (B), or by two-sided Student’s t-test with Welch’s correction (A). See the gating strategy for the flow cytometry analysis in fig. S8A (A) and B (B), and the representative density plots in fig. S8C (A) and D (B). (**C** and **D**) AMPK is required for aldometanib-induced infiltration of CD8^+^ T cells into HCC tissues. HCC tissues from the wildtype or *AMPKα*-LKO DEN-HFD mice (**C**), or from xenografts derived from wildtype or *AMPKα*^-/-^ Hepa1-6 cells transplanted into wildtype or *AMPKα*-LKO mice (D), were subjected to immunohistochemistry staining for CD8a. Representative images are shown on the left panels, and the percentages of CD8a-positive areas within the tumor regions were calculated and are shown on the right panel (means ± s.e.m., *n* represents the number of mice, and are labeled in each panel; *P* values were calculated by two-way ANOVA, followed by Tukey). The scale bars are 100 μm. (**E** to **H**) Tumouricidal activities are detected in the CD8^+^ T cells-positive areas of HCC. HCC tissues from DEN-HFD mice ((E) and (F)) or Hepa1-6-derived xenografts ((G) and (H); using the Hepa1-6-non-transplated mouse liver, or naïve mouse liver, as a control) were digested and stained with CD8-Alexa Fluor 488 ((E) and (F); as in (A)) or CD8-APC ((G) and (H); as in (B)). Cells were further stained with granzyme B-PE ((E) and (G)) or IFNγ-PE ((F) and (H)) antibody to differentiate the sub-population of CD8^+^ T cells with high tumouricidal activity by flow cytometry analysis. Statistical analysis data are shown as violin plots; *n* = 18 samples from 6 mice ((E) and (F)), or represents the number of mice, and are labelled in each panel ((G) and (H)); *P* values calculated by two-sided Student’s *t*-test ((E) to (H)). See the gating strategy for the flow cytometry analysis in fig. S8A ((E) and (F)), B ((G) and (H)), and the representative density plots in fig. S8E (E), G (F), F(G), and H (H). (**I** to **L**) Aldometanib eases the immune barriers in the HCC microenvironment. HCC tissues from Hepa1-6-derived xenografts were digested as in (B). Cells were then stained with Ly6G-Alexa Fluor 488, CD45-PerCP-Cy5.5, and CD11b-APC antibodies to label neutrophils (I), the CD11b-APC and the F4/80-PE antibodies to label macrophages (J), the CD11c-PerCP-Cy5.5, CD11b-APC, and CD45-Alexa Fluor 488 antibodies to label lymphoid DCs (K), and the MHC II-PE, CD11c-PerCP-Cy5.5, and CD11b-APC antibodies to label the MHC II-expressed myeloid DCs (L). Statistical analyses are shown as violin plots; *n* = 18 samples collected from 6 mice, with *P* values calculated by two-sided Student’s *t*-test (L), or by two-sided Student’s *t*-test with Welch’s correction ((I) to (K)). See the gating strategy for the flow cytometry analysis in fig. S8I ((I) and (J)), 8L ((K) and (L)), and the representative density plots in fig. S8J (I), K (J), M (K), and N (L). **(M)** Depletion of CD8^+^ T cells impedes aldometanib-induced suppression of HCC. Wildtype C57BL/6 mice were transplanted with wildtype Hepa1-6 cells, followed by treatment with aldometanib starting from day 2 post-transplantation (depicted in the upper panel). Starting from day 6 after the transplantation, the mice were intravenously injected with neutralizing antibodies against CD8^+^ T cells twice a week. HCC tissue samples were collected on day 17 post-transplantation, followed by the determination of tumor weights (means ± s.e.m., n represents the number of mice, and are labeled in each panel, with *P* values calculated by two-way ANOVA, followed by Tukey). See also the morphologies of HCC in fig. S9F (using H&E staining) and the validation data for the effectiveness of CD8^+^ T cell depletion in fig. S9G (using immunohistochemistry staining). **(N)** Re-introduction of CD8^+^ T cells restores the ability of Aldometanib to inhibit HCC in immunodeficient NOD-SCID mice. The NOD-SCID mice were intraperitoneally injected with human PBMCs. At 3 days after the injection, mice were transplanted with human HCC cells into the left lobe of the liver. HCC tissue samples were collected on day 17, followed by the determination of tumor weights (means ± s.e.m., n represents the number of mice, and are labeled in each panel, with *P* values calculated by two-way ANOVA, followed by Tukey). See also tumor:body weight ratios and the infiltration of CD8^+^ T cells in fig. S10, M to R. Experiments in this figure were performed three times.

Compared to CD8^+^ T cells, we found that aldometanib did not significantly promote the infiltration of other tumor-infiltrating lymphocytes (TIL), including the Th17 cells (fig. S8Q) and NK cells (CD3-negative and NK1.1-positive, or CD3^-^NK1.1^+^; fig. S11A and B). Although aldometanib was able to help recruit NKT cells (CD3^+^NK1.1^+^) into HCC tissues, the contents of the NKT cells remained lower than that in the livers of healthy mice (fig. S11, A and B). We also did not observe any change in eosinophils and basophils before or after aldometanib treatment (fig. S11, C and D). The data above indicate that aldometanib stimulates the recruitment of CD8^+^ T cells into the HCC tumors to exert tumouricidal activity.

## Discussion

We have shown that the non-cytotoxic glucose starvation mimetic aldometanib treatment can reduce the sizes of tumors in both of the HCC spontaneously developed in DEN-HFD and in *MYC*;*Trp53*^−/−^ mice, and xenografts of various origins, including Hepa1-6 and patient-derived HCC cells. Remarkably, aldometanib treatment allows the tumor-carrying mice to live to normal ages, as determined in the DEN-HFD mice. We have also revealed that AMPK in the para-tumor tissue plays a dominant role as assessed in xenografts, to such extent that even AMPK-null HCC can be as effectively suppressed by aldometanib, so long as the AMPK in the host tissue is present. This makes it unlikely that aldometanib-induced AMPK activation within the tumor cells helps suppress the tumor growth or cleanse the tumor cells. It is also worth pointing out that the mild inhibitory effect of aldometanib on the proliferation of various HCC cells, possibly through reduced levels of anabolic activities after AMPK activation, does not account for the reduction of tumor sizes, for the xenografts were formed prior to aldometanib administration.

We have also shown that while aldometanib does not possess cytotoxicity towards cancerous cells or normal cells, it robustly induces tumor cells to undergo apoptosis and necroptosis, signs of tumouricidal activity, which depends on the infiltration of the cytotoxic lymphocytes CD8^+^ T cells. As AMPK is known to play a central role in both anabolism and catabolism, influencing levels/functions of both metabolites and proteins, it is reasonable to suggest that aldometanib, via activation of AMPK, “clears” the way for the mobilization of immune cells, particularly the CD8^+^ T cells, to induce the cytotoxicity on the tumor tissues. To explore what accompanies the promotion of the migration of CD8^+^ T cells, we analyzed chemokines such as CXCL9, CXCL10, and CXCL11 that are required for the infiltration of CD8^+^ T cells (*62–64*). Surprisingly, these chemokines remained unchanged, or even decreased after aldometanib treatment, both in tumor and para-tumor tissues (fig. S12, A to F and fig. S13, A to I). As a control, lipopolysaccharide (LPS) significantly increased the levels of these cytokines in the liver (fig. S13J). It is possible that aldometanib, by rebalancing metabolism to improve the microenvironment within the host tissues and/or the microenvironment between the tumor tissue and para-tumor tissue, does not rely on these cytokines to promote the infiltration of CD8^+^ T cells into HCC. It remains a conundrum what AMPK activation does to improve the balance of metabolites and likely the abundance and properties of proteins to remove the immune barrier upon the activation by aldometanib.

Current immunotherapies for HCC are often restricted to certain etiologies or subtypes of HCC (*65*), because these therapies must be tailored to restore the particular types of immune cells that are dysregulated in a specific HCC etiology, in order to counteract the immunosuppression. For example, NASH-induced HCCs often exhibit increased levels of IL-10, one of the cytokines that can suppress CD8^+^ T cells, which is secreted by the circulating IgA^+^ plasmocytes (*66*). In hepatitis B virus-induced HCC, IL-10 is mainly produced by intrahepatic Breg cells (*67, 68*), while in hepatitis C virus-induced HCC, it is mainly from the circulating CD4^+^CD25^+^ Treg cells (*69*). In addition, in NASH-induced HCCs, a specific subset of T cells, known as PD-1^+^CXCR6^+^CD8^+^ T cells, plays a crucial role in creating an immunosuppressive TME and hindering the response to immune checkpoint inhibitors (*70, 71*). In this study, we have studied HCCs of DEN-HFD mice and *MYC*;*Trp53*^−/−^ mice, and xenografts of various origins, including Hepa1-6 and patient-derived HCC cells. We have shown that aldometanib can elicit inhibitory effects on all the types of HCC tested, depending on AMPK. It is possible that AMPK activation by aldometanib may improve the overall metabolic network to clear the way for CD8^+^ T cells to migrate and reach various types of HCC. Consistent with this notion, we and others have shown that caloric restriction (CR) or metformin, which can activate AMPK in the liver (*72, 73*), showed some effects on the progression of HCC in mice ((*37–40, 74, 75*) and fig. S14, A to H). Also of some relevance, metformin was shown to enhance the efficacy of immunotherapies (*76*). In short, we have demonstrated that metabolic intervention, as exemplified by the application of aldometanib, can allow cancers to become a manageable disease, enabling cancer patients to live with cancer.

## ACKNOWLEDGMENTS

We thank Dr. Sean Morrison (University of Texas Southwestern Medical Center) for providing the *AMPKα1*^F/F^ (The Jackson Laboratory, 014141), and *AMPKα2*^F/F^ (The Jackson Laboratory, 014142) mice; Su-Qin Wu, Ying He and Jing Song (Laboratory Animal Research Centre, Xiamen University) for mouse in vitro fertilization; Cixiong Zhang and Zicheng Huang (Core Facility of Biomedical Sciences, Xiamen University) for HPLC-MS and MRI analysis, respectively; Zhenning Wang and Xuanzhang Huang (Department of Surgical Oncology and General Surgery, the First Hospital of China Medical University) for analyzing the RNA sequencing data; Lintao Song (School of Pharmaceutical Sciences, Wenzhou Medical University) for assistance with HCC induction and analysis in mice; Yanping Chen, Jianchao Wang, Dongmei Zhou, Jianping Lu, Shuyi Lu, Weifeng Zhu, and Menghan Fang (Clinical Oncology School of Fujian Medical University, Fujian Cancer Hospital) for the analysis of the pathological features of patient HCC samples; Zhiyun Ye, Wei Hong and Tian-Yu Tang for secretarial assistance; and all the other members of the S.-C.L. laboratory for various technical assistance.

## Author contributions

H.-H.H., C.-S.Z., and S.-C.L. conceived the study and designed the experiments. H.-H.H. determined the effects of aldometanib on HCC suppression with the assistance of M.L., J.C., D.C., and S.L. X.W. and B.L. determined the effects of aldometanib on CD8^+^ T infiltration, with the assistance of H.C., H.W., F.C. and G.C. X.W. and B.L. isolated the patient-derived HCC cells and generated the patient-derived HCC xenograft model under the guidance of F.C. and J.Z. B.L. established the mouse primary HCC cell line. J.W. bred the mouse strains. C.-S.Z. and M.L. analyzed the lifespan of HCC-bearing mice. M.Y. and Z.W. synthesized aldometanib and prepared the preparations of aldometanib for mouse studies, under the guidance of X.D. G.C. and Y.C. analyzed the pathological features of patient HCC tissues. Z.-Z.Z. performed the RNA-sequencing analysis. J.L. and S.-C.L. supervised the project. C.-S.Z. and S.-C.L. wrote the manuscript.

## Data and materials availability

The data supporting the findings of this study are available within the paper and its Supplementary material files. The raw RNA sequencing data corresponding to the expression of chemokines in the mouse tissues have been deposited in the Genome Sequence Archive (*77*) in National Genomics Data Center, China National Center for Bioinformation/Beijing Institute of Genomics, Chinese Academy of Sciences (GSA: CRA019674 and CRA019771) that are publicly accessible at https://ngdc.cncb.ac.cn/gsa. Materials and reagents are available upon request. Questions regarding the details of experiments are welcome. Full immunoblots are provided as “Full scans”. Source data for statistical analyses are provided with this paper. The analysis was performed using standard protocols with previously described computational tools. No custom code was used in this study.

## SUPPLEMENTARY MATERIALS

Materials and Methods Figs. S1 to S14

Table S1

References (1–106)

