## Supplementary Materials for "AMPK acts to remove immune barriers to CD8^+^ T cell-mediated immunity against hepatocellular carcinoma"

**The PDF file includes:**

Materials and Methods
Figs. S1 to S14
Table S1
References

#### Materials and Methods

Data reporting

The chosen sample sizes were similar to those used in this field: *n* = 4-8 samples were used to evaluate the levels of metabolites in serum (*35, 41*) and tissues (*30, 35, 41*); *n* = 4-15 samples to determine the mRNA levels of a specific gene (*29, 41, 43*); *n* = 4-6 samples to determine the expression levels and phosphorylation levels of a specific protein (*20, 29, 35*); *n* = 6-15 mice to induce HCC (*20, 42, 43, 78, 79*); *n* = 6 mice to determine of hepatic TAG (*26, 35, 41*); *n* = 3-5 mice to determine the formation of HCC using MRI (*70, 80, 81*); *n* = 4-18 samples for the flow cytometry analysis of immune cells (*78, 79*); *n* = 3-14 mice for IHC and H&E staining (*20, 41, 78, 79*); *n* = 33 mice for generating survival curves of tumor-bearing mice (*20, 42, 43*); *n* = 6-15 mice for determination of serum ALT and AST (*41, 43*). No statistical methods were used to predetermine the sample size. All experimental findings were repeated as stated in the figure legends, and all additional replication attempts were successful. For animal experiments, mice were housed under the same condition or place. For cell experiments, cells of each genotype were cultured in the same condition and were seeded in parallel for different treatments. Each experiment was designed and performed along with proper controls, and samples for comparison were collected and analyzed under the same conditions. Randomization was applied wherever possible. For example, during MS analyses (metabolites and pharmacokinetics), samples were processed and subjected to the mass spectrometer in random orders. For animal experiments, age-matched litter-mate animals in each genotype were randomly assigned to aldometanib treatments. In cell experiments, cells were randomly assigned to different treatments. Similarly, during microscopy data collection and statistical analyses, the fields of view were chosen on a random basis, and were often performed by different operators, preventing potentially biased selection for desired phenotypes. Otherwise, randomization was not performed. For example, when performing immunoblotting, samples needed to be loaded in a specific order to generate the final figures. Blinding was applied wherever possible. For example, samples or cages during sample collection and processing were labelled as code names that were later revealed by the individual who picked and treated animals or cells, but did not participate in sample collection and processing, until assessing outcome.

Mouse strains

Protocols for all rodent experiments were approved by the Institutional Animal Care and the Animal Committee of Xiamen University (XMULAC20180028 and XMULAC20220050). Unless stated otherwise, mice were housed with free access to water and standard diet (65% carbohydrate, 11% fat, 24% protein) under specific pathogen-free conditions. The light was on from 8:00 to 20:00, with the temperature kept at 21-24 °C and humidity at 40-70%. Only male mice were used in the study, and male littermate controls were used throughout the study.

For CR, mice were individually caged for 1 week before the treatment. Each mouse was fed with 2.5 g of standard diet (approximately 70% of ad libitum food intake for a mouse at 8 weeks old and older) at 5 p.m. each day.

Wildtype C57BL/6J (#000664) and BALB/c (#000651) mice were obtained from The Jackson Laboratory, and the BALB/c nude mice (#401) and NOD-SCID mice (#394) from Charles River Beijing branch (Vitalriver). *AMPKα1*$^{F/F}$ (#014141) and *AMPKα2*$^{F/F}$ mice (#014142) were obtained from Jackson Laboratory, provided by Dr. Sean Morrison. *AMPKα*-LKO mice were generated by crossing *AMPKα1/2*$^{F/F}$ mice with *Alb-Cre* mice, as described previously (*41*). The primers used for genotyping the *AMPKα*-LKO mice were as follows: 5'-CCCACCATCACTCCATCTCT-3' and 5'-AGCCTGCTTGGCACACTTAT-3' for *AMPKα1*, 5'-GCAGGCGAATTTCTGAGTTC-3' and 5'-TCCCCTTGAACAAGCATACC-3' for *AMPKα2*, and 5'-ATGAAATGCGAGGTAAGTATGG-3' and 5'-CGCCGCATAACCAGTGAAAC-3' for *Alb-Cre*.

In this study, mice of following ages were used: a) for generating HCC xenograft-bearing mice: wildtype or AMPKα-LKO C57BL/6J mice of 8 weeks old (for transplanting Hepa1-6 cells), NOD-SCID mice of 8 weeks old (for transplanting human patient-derived HCC cells), or BALB/c nude mice of 8 weeks old (for transplanting Huh7 and JHH-7 cells); b) to induce DEN-HFD HCC: wildtype or AMPKα-LKO C57BL/6J mice of 4 weeks old; c) to induce *MYC*;*Trp53*$^{-/-}$ HCC: wildtype BALB/c mice of 12 weeks old; d) to analyze AMPK activation, adenylate levels, and hepatoma phenotypes in liver tissues: DEN-HCC mice of 41 weeks old (treated with either vehicle or aldometanib dissolved in drinking water for 1 week, starting at 40 weeks old), Hepa1-6-derived xenograft-bearing mice of 10 weeks old (also treated with either vehicle or aldometanib for 1 week, starting from the day 2 after cell transplantation), or *MYC*;*Trp53*$^{-/-}$ mice of 16 weeks old (treated similarly starting from 15 weeks old); e) to determine the concentrations of aldometanib in serum and liver tissues: DEN-HCC mice of 41 weeks old (treated as above) and Hepa1-6-derived xenograft-bearing mice of 10 weeks old (treated as above); f) to assess the composition of immune cells in HCC tissues: Hepa1-6-derived xenograft-bearing mice of 10 weeks old (treated with either vehicle or aldometanib for a total of 8 days, starting from the day 2 after cell transplantation) and DEN-HFD mice of 40 weeks old (treated for 8 weeks starting from 32 weeks old); and g) to analyze chemokine levels using RT-PCR or RNA sequencing: Hepa1-6-derived xenograft-bearing mice of 10 weeks old (treated for a total of 8 days, starting from the day 2 after cell transplantation), DEN-HFD mice of 40 weeks old (treated for 8 weeks starting from 32 weeks old), wildtype C57BL/6J mice of 8 weeks old, and wildtype mice of 4 weeks old were used to isolate primary hepatocytes.

<u>Formulation of Aldometanib</u>
Aldometanib was formulated as described previously (*41*). Briefly, for cell-based experiments, aldometanib powder was dissolved in DMSO, and was aliquoted and stored at 4 °C. The solution was incubated in a 37 °C water bath for 10 min (until no precipitate was visible) before adding to the culture medium. For mouse experiments, aldometanib was formulated in 10 % (w/v) Kolliphor HS 15 before the experiment. Briefly, to prepare 3 l of 1 mg/mL Aldometanib stock solution, 3 g of aldometanib was dissolved in 300 mL of ethanol, followed by mixing with 300 g of Kolliphor HS 15. The mixture was stirred at room temperature until Kolliphor HS 15 was completely dissolved. The ethanol in the solution was then evaporated on a rotary evaporator (Rotavapor R-300, BUCHI) at 90 rpm, 45 °C, followed by mixing with 2,700 mL of water. The control vehicle was similarly prepared, with no aldometanib added. Both the Aldometanib solution and the vehicle were stored at room temperature, and were used within 1 month after the preparation.

<u>Isolation of patient-derived HCC cells</u>
Patient HCC tissues were obtained from the Fujian Cancer Hospital. All procedures were conducted in accordance with the Declaration of Helsinki and were approved by the Tumor Ethics Committee of Fujian (K2023-003-01). All participants signed an informed consent form before enrolment. The collected HCC tissues showed a macrotrabecular-massive type and tested positive

for cirrhosis. Determined by immunohistochemistry staining, the HCC tissues also tested positive for Arg-1, CD34 (only vascular expression), and GS, but negative for HepPar-1, CK7, CK19, and CD10. In addition, approximately 30% of the nuclei were positively labeled for Ki-67, with partial positivity for HSP70 and Glypican-3.

The patient HCC tissues were collected and maintained in NOD-SCID mice as described previously (*82-84*), with minor modifications. Briefly, freshly excised HCC tissues were placed in serum-free DMEM in an ice bath, and sliced to a diameter of 1-2 mm at the surgical site. After washing with ice-cold serum-free DMEM 2 times, tissues were subcutaneously implanted into the right flanks of 8-week-old NOD-SCID mice. The tumor tissues were collected and sliced into 1 mm diameter after reaching approximately 1,000 $mm^3$ in the NOD-SCID mice, and then digested with 10 mL of type I collagenase (1 mg/mL final concentration, in serum-free DMEM) in a petri dish at 37 °C, 60 rpm for 10 min on an orbital shaker. After passing the digested tissue through a 70-μm cell strainer (cat. 352350; BD Falcon), cells were centrifuged in 400*g* for 5 min at room temperature, followed by suspension with serum-free DMEM and then transplanted into the liver of NOD-SCID mice as described below in "HCC-HFD, *MYC*;*Trp53*$^{-/-}$ mice and HCC xenografts".

HCC-HFD, *MYC*;*Trp53*$^{-/-}$ mice and HCC xenografts

The DEN-HFD mouse HCC model was established as described previously (*43*), with minor modifications. Briefly, C57BL/6J mice were given two intraperitoneal injections of 80 mg/kg DEN, one on postnatal day 28 and the other on postnatal day 35. Two weeks after the second injection, the mice were switched to HFD (60% calories from fat; D12492, Research Diets) for another 41 weeks (except for the analysis of AMPK activation and AMP levels in the HCC tissues, the duration of HFD feeding was 34 weeks, as shown in fig. S1, A and B). Mice were then administered aldometanib in the drinking water. At the end of the study, the mice were euthanized, and their liver tissues were collected for analysis of the diameters of tumors using Vernier calipers (cat. 150T, Meinaite tools, China).

The *MYC*;*Trp53*$^{-/-}$ HCC mice were established as described previously (*42, 79*), with minor modifications. Briefly, BALB/c mice of 12 weeks old were hydrodynamically injected with a plasmid mixture consisting of 30 μg of pX330-sg-p53 (ref. (*85*); #59910, Addgene, gift from Dr. Tyler Jacks), 30 μg of PT3-EF1a-Myc (#92046, Addgene, gift from Dr. Xin Chen), and 7.5 μg (4:1 ratio) of pCMV/SB10 transposase-encoding plasmids (#24551, Addgene, gift from Dr. Perry Hackett), all freshly dissolved in 2 mL of 0.9% NaCl (w/v) solution, through the tail vein. For each mouse, a total volume of mixture corresponding to 10% of body weight was injected into the lateral tail vein within 7 s. Mice were then intraperitoneally injected with 10% (v/v in corn oil) $CCl_4$ at a dose of 5 mL/kg twice per week, starting at 1 week after hydrodynamic tail-vein injection for 3 weeks.

The Hepa1-6-derived hepatoma xenografts (orthotopic implantation) were established in situ as described previously (*86-88*), with minor modifications. Briefly, the Hepa1-6 cells were trypsinized, washed with serum-free DMEM medium, and then resuspended in serum-free DMEM medium. Some $1.5 \times 10^6$ wildtype Hepa1-6 cells or $2 \times 10^6$ *AMPKα*$^{-/-}$ Hepa1-6 cells, suspended in 40 μL of serum-free DMEM medium, were injected slowly into the left liver lobe of each C57BL/6J mouse of 8 weeks old. After the injection, the liver surface at the needle site was gently covered with a sterile cotton swab for around 2 min to minimize bleeding and potential backflow/leakage. The mice were then administered with aldometanib in the drinking water as described in Fig. 1F, followed by analysis of tumor sizes at indicated time points. Hepatoma xenografts derived from human HCC cell lines (JHH-7 and Huh7) were similarly established, except that $1 \times 10^7$ cells were transplanted into 8-week-old BALB/c nude mice. The mice were then administered with aldometanib in the drinking water as described in fig. 10, A and G, followed by the analysis of tumor sizes at indicated time points.

To deplete the $CD8^+$ T cells in the Hepa1-6-derived xenograft-bearing mice, 200 μg of neutralizing antibody against CD8a was intravenously injected into the mice twice a week, starting from day 6 after cell transplantation, followed by treatment with aldometanib as in Fig. 4M. The efficiency of $CD8^+$ T cell depletion was confirmed by immunohistochemistry at day 17 after cell transplantation.

The patient-derived HCC xenografts were established as described previously (*89-91*), with minor modifications. Briefly, 8-week-old NOD-SCID mice were injected each with $2 \times 10^7$ PBMCs that were freshly prepared from the whole blood of healthy human donors. The preparation of PBMCs was performed by mixing the blood with PBS in a 1:5 ratio, and then 5 mL of the mixture containing PBMCs was transferred onto the upper layer of 3 mL of ficoll, followed by centrifuge for 20 min at 2,000*g* at 25 °C. Note that the centrifuge was set at minimum acceleration and deactivated the brake program, as it would otherwise disturb the ficoll gradient and, in turn, lower the cell yield. After centrifugation, approximately 1 mL of the fraction located above the ficoll surface, which contained the PBMCs, was collected using a Pasteur pipette. This fraction was mixed with 10 mL of PBS and centrifuged again for 5 min at 400*g* and 25 °C. At 3 days after the PBMC injection, $1 \times 10^7$ human HCC cells, suspended in 40 μL of serum-free DMEM, were transplanted into the left lower lobe of the liver of the mice. Mice were then treated with aldometanib in the drinking water, as in Fig. 4N.

Evaluation of the lifespan of HCC-bearing mice

The survival lifespans of DEN-HFD mice were determined according to previous reports (*41, 43*), with minor modifications. Briefly, mice were examined every morning for signs of illness, and the severely moribund mice were censored. A mouse was considered severely moribund if it showed more than one of the following clinical signs: a) inability to eat or to drink; b) severe lethargy, as indicated by lack of response such as a reluctance to move when gently prodded with a blunt-tip tweezer; or c) severe balance instability or gait disturbance. Mice found dead were also recorded at each daily inspection. Aldometanib was administered starting at 12 weeks of age and continued throughout the lifespan of the mice.

In the cohort shown in Fig. 1A and table S1, the experiment started with a total of 104 male mice: 53 in the vehicle group and 51 with the administration of aldometanib. Along the experiment, removed (censored) from the study were 38 mice (20, vehicle; 18, aldometanib). The reasons for removal included fighting (6, vehicle; 3, aldometanib), paralysis (loss of walking ability: 5, vehicle; 6, aldometanib), severe ulcerative dermatitis (5, vehicle; 4, aldometanib), and symptoms of gnawing or bruxing (the presence of long, spiral incisors preventing the mouse from eating: 4, vehicle; 5, aldometanib). Such censored mice were not included in the calculation of lifespans.

Magnetic resonance imaging

To detect the local growth of hepatoma in DEN-HFD mouse, liver magnetic resonance imaging

(MRI) was performed using a 9.4-T magnetic resonance scanner (Biospec 94/20 USR, Bruker) as described previously (*81, 92*). Briefly, the mouse was anaesthetized and placed in a prone position with its extremities extended on the imaging bed. After positioning the mouse coil at the iso-center of the scanner, imaging was performed using a T2-weighted rapid acquisition with relaxation enhancement (RARE) sequence. The parameters were as follows: a) echo times: 33 ms; b) repetition time: 2,500 ms; c) averages: 2; d) repetitions: 1; e) echo spacing: 11 ms; f) rare factor: 8; g) slices: 20; h) slice orientation: axial; h) slice thickness and distance: 1 mm; i) image size: 256 × 256; j) field of view: 35 × 35 mm; k) echo images: 1; l) excitation angle: 90°; m) refocusing angle: 180°; n) dummy scans: 2; o) dummy duration: 5,000 ms; p) motion averaging, flip back, and fat suppression: ticked; q) dimension: 2D; r) isotropic: off; s) anti aliasing: 1 × 1; t) object ordering mode: interlaced; u) read offset: 2.023 mm; v) phase offset: -2.262 mm; w) slice offset: 0.792 mm; x) slice gap mode: non-contiguous; y) bandwidth: 36764.7 Hz; z) excitation pulse and refocusing pulse: calculated; and aa) auto repetition spoiler and auto echo spoiler: ticked.

Cell lines

Throughout this study, no cell line used is listed as misidentified by the International Cell Line Authentication Committee (https://iclac.org/databases/cross-contaminations/). HEK293T (cat. CRL-3216), Hepa1-6 (cat. CRL-1830), and BNL (cat. TIB-73) cells were purchased from ATCC, Huh7 cells (cat. CL-0120) from Procell, and JHH-7 (cat. CTCC-004-0053) from Meisen CTCC. All cells were verified to be free of mycoplasma contamination and authenticated by STR sequencing. All of the cell lines were maintained in DMEM supplemented with 10% FBS, 100 IU penicillin, and 0.15 mg/mL streptomycin, except that JHH-7 cells in DMEM/F-12 medium supplemented with 10% FBS, 100 IU penicillin, 0.15 mg/mL streptomycin. Cells were cultured at 37 °C in a humidified incubator containing 5% $CO_2$.

The genes for *PRKAA1* and *PRKAA2* were deleted from Hepa1-6 cells using the CRISPR-Cas9 system. The sgRNAs were designed as described previously (*93*). Nucleotides were annealed to complements containing the cloning tag aaac, and inserted into the back-to-back *Bsm*B I restriction sites of the lentiCRISPRv2 vector (#52961, Addgene). The sequences of sgRNAs are as follows: 5'- GGGCCGCAATAAAGATATC-3' for mouse *PRKAA1*; and 5'-GGGAGCCCGTGCGCCGAACA-3' for mouse *PRKAA2*. The constructs were then subjected to lentivirus packaging using HEK293T cells that were transfected with 1.5 μg of DNA in Lipofectamine 2000 transfection reagent per well of a 6-well plate. At 30 h after transfection, the virus (DMEM supplemented with 10% FBS, MEM non-essential amino acids, and sodium pyruvate; approximately 2 mL) was collected and centrifuged at 5,000*g* for 3 min at room temperature. The supernatant was mixed with 10 μg/mL (final concentration) polybrene; the virus mixture was then added to Hepa1-6 cells of 15% confluence, followed by centrifuging at 3,000*g* for 30 min at room temperature (spinfection). Cells were then incubated with the virus for another 48 h before refreshing the medium. Cells of near confluence were single-cell-sorted into 96-well dishes. The resultant clones were expanded and evaluated for knockout status by sequencing.

Primary mouse hepatocytes and primary HCC cells

Primary mouse hepatocytes were isolated with a modified two-step perfusion method using Liver Perfusion Medium and Liver Digest Medium as described previously (*29*). Briefly, before isolation of hepatocytes, mice were first anaesthetized, followed by insertion of a 0.72 mm × 19 mm intravenous catheter into postcava. After cutting off the portal vein, mice were perfused with 50 mL of Liver Perfusion Medium at a rate of 5 mL/min, followed with 50 mL of Liver Digest Medium at a rate of 2.5 mL/min. The digested liver was then briefly rinsed by PBS, and then dismembered by gently tearing apart the Glisson's capsule with two sterilized, needle-pointed tweezers on a 6-cm dish containing 3 mL of PBS. The dispersed cells were mixed with 10 mL of ice-cold William's medium E plus 10% FBS, and were filtered by passing through a 100-μm cell strainer (cat. 352360, BD Falcon). Cells were then centrifuged at 50*g* at 4 °C for 2 min, followed by washing twice with 10 mL of ice-cold William's medium E plus 10% FBS. Cells were then immediately plated (at 60-70% confluence) in collagen-coated six-well plates in William's medium E plus 10% FBS and GlutaMAX, 100 IU penicillin and 0.1 mg/mL streptomycin, and were maintained at 37 °C in a humidified incubator containing 5% $CO_2$. After 2 h of attachment, the medium was replaced with fresh William's medium E with 1% (w/v) BSA and GlutaMAX for another 12 h before further use.

The mouse primary HCC cells were isolated from the DEN/$CCl_4$-induced HCC tissues (generated as described previously (*94*), with minor modifications (*79*)), and were cultured in vitro as described previously (*95, 96*). Briefly, C57BL/6J mice at 4 weeks old and 5 weeks old were intraperitoneally injected with 80 mg/kg DEN twice, followed by intraperitoneal injection with 10% (v/v in corn oil) $CCl_4$ at a dose of 5 mL/kg every week from 8 weeks old. Tumor tissues were collected from mice at 40 weeks old and were sliced into cubes with a diameter of 1-2 mm in serum-free DMEM in an ice bath. Tumor tissues were washed with ice-cold serum-free DMEM 2 times, and then intraperitoneally implanted into the 8-week-old C57BL/6J mice. After 1 month, the tumor tissues were collected, minced, and digested with 10 mL of type I collagenase (1 mg/mL final concentration, in serum-free DMEM) in a petri dish at 37 °C for 10 min. The cells were then centrifuged at 400*g* for 5 min at room temperature, suspended, and cultured in DMEM medium supplemented with 10% FBS and 10 μM Y-27632 for 24 h. The medium was then refreshed to DMEM medium supplemented with 10% FBS every two days. Cells were ready for further uses after at least 3 passages of culture.

<u>Reagents</u>
Rabbit anti-phospho-AMPKα-Thr172 (cat. #2535, RRID: AB_331250; 1:1,000 dilution for IB), anti-AMPKα (cat. #2532, RRID: AB_330331; 1:1,000 for IB), anti-phospho-ACC-Ser79 (cat. #3661, RRID: AB_330337; 1:1,000 for IB), anti-ACC (cat. #3662, RRID: AB_2219400; 1:1,000 for IB), anti-phospho-p70 S6K-Thr389 (cat. #9234, RRID: AB_2269803; 1:1,000 for IB), anti-p70 S6K (cat. #2708, RRID: AB_390722; 1:1,000 for IB), anti-PUMA (cat. #98672, RRID: AB_3096180; 1:1,000 for IB), anti-caspase-3 (cat. #14220, RRID: AB_2798429; 1:1,000 for IB), anti-caspase-7 (cat. #12827, RRID: AB_2687912; 1:1,000 for IB), anti-GSDMD (cat. #39754, RRID: AB_2916333; 1:1,000 for IB), anti-Ki-67 (cat. #12202, RRID: AB_2620142; 1: 200 for IHC), and mouse anti-caspase-9 (cat. #9508, RRID: AB_2068620; 1:1,000 for IB) antibodies were purchased from Cell Signaling Technology. Rabbit anti-tubulin (cat. #10068-1-AP, RRID: AB_2303998; 1:1,000 for IB), anti-GSDME (cat. #13075-1-AP, RRID: AB_2093053; 1:1,000 for IB), anti-AFP (cat. #14550-1-AP, RRID: AB_2223933; 1: 100 for IHC), anti-CXCL9 (cat. #22355-1-AP, RRID: AB_2879086; 1: 500 for IHC), and anti-CXCL10 (cat. #10937-1-AP, RRID: AB_2088002; 1: 200 for IHC) antibodies were purchased from Proteintech. Rabbit anti-CXCL11 (cat. #orb13425, RRID: AB_10749546; 1: 100 for IHC) was purchased from Biorbyt. Rat anti-CD16/CD32 (cat. #14-0161-86, RRID: AB_467135; 1: 100 for F) antibody was purchased from Invitrogen. Rat anti-IFNγ PE-CF594 (cat. #562303, RRID: AB_11153140; 1: 200 for F) antibody

was purchased from BD Biosciences. Rat anti-CD45 Alexa Fluor 488 (cat. #53-0451-82, RRID: AB_2848416); anti-CD45 PerCP-Cyanine5.5 (PerCP-Cy5.5) (cat. #45-0451-82, RRID: AB_1107002; 1: 200 for F), anti-CD8a APC (cat. #17-0081-82, RRID: AB_469335; 1: 200 for F), anti-CD11b APC (cat. #17-0112-83, RRID: AB_469344; 1: 200 for F), anti-CD11c PerCP-Cy5.5 (cat. #45-0114-82, RRID: AB_925727; 1: 200 for F), anti-granzyme B PE (cat. #12-8898-82, RRID: AB_10870787; 1: 200 for F), anti-IL-1β PE (cat. #12-7114-82, RRID: AB_10732630; 1: 200 for F), anti-IL-4 PE (cat. #12-7041-82, RRID: AB_466156; 1: 200 for F), anti-IL-17a PE (cat. #12-7177-81, RRID: AB_763582; 1: 200 for F), and mouse anti-NK1.1 APC (cat. #17-5941-82, RRID: AB_469479; 1: 200 for F) antibodies were purchased from eBioscience. Rat anti-CD3 PE (cat. #100206, RRID: AB_312663; 1: 200 for F), anti-CD4 Alexa Fluor 488 (cat. #100529, RRID: AB_389303; 1: 200 for F), anti-CD8a Alexa Fluor 488 (cat. # 100723, RRID: AB_389304; 1: 200 for F), anti-F4/80 PE (cat. # 123110, RRID: AB_893486; 1: 200 for F), anti-Ly6G-1A8 Alexa Fluor 488 (cat. # 127626, RRID: AB_2561339; 1: 200 for F), anti-CD4 APC (cat. #116014, RRID: AB_2563024; 1: 200 for F), anti-CD19 PerCP-Cy5.5 (cat. #152406, RRID: AB_2629815; 1: 200 for F), anti-CD206 Alexa Fluor 488 (cat. #141710, RRID: AB_10900445; 1: 200 for F), anti-MHC II PE (cat. #107608, RRID: AB_313323; 1: 200 for F), anti-CD170-Siglec-F PE (cat. #155506, RRID: AB_2750234; 1: 200 for F), CCR3-APC (cat. #144511, RRID: AB_2565737; 1: 200 for F), and Armenian hamster anti-TCRγ/δ APC (cat. #118116, RRID: AB_1731813; 1: 200 for F) and anti-FcεRIα Alexa Fluor 488 (cat. #134330, RRID: AB_2687239; 1: 200 for F) antibodies were purchased from BioLegend. The neutralizing antibody of $CD8^+$ T cells, rat anti-CD8a (clone 2.43; cat. BE0061, RRID: AB_1125541), and its isotype control antibody (rat IgG2b isotype control; clone LTF-2; cat. #BE0090, RRID: AB_1107780) were purchased from BioXCell. Rat anti-CD8 (cat. #ab217344, RRID: AB_2890649; 1: 2,000 for IHC of mouse CD8a), anti-NK1.1 (cat. #ab289542, RRID: AB_3094493; 1: 100 for IHC), rabbit anti-CD8 (cat. #ab245118, RRID: AB_3068617; 1: 2,000 for IHC of human CD8a), and mouse anti-Ly6G (cat. #ab210204, 1: 100 for IHC) were purchased from Abcam. The horseradish peroxidase (HRP)-conjugated goat anti-mouse IgG (cat. #115-035-003, RRID: AB_10015289; 1: 5,000 for IB) and goat anti-rabbit IgG (cat. #111-035-003, RRID: AB_2313567; 1: 5,000 for IB) antibodies were purchased from Jackson ImmunoResearch.

Aldometanib was synthesized as described previously (*41*), and is now available at MedChemExpress (cat. HY-148189), GLPBIO (cat. GC66024), and CymitQuimica (cat. TM-T60122). DMSO (cat. D2650), methanol (cat. 646377), ethanol (cat. 459836), isopropanol (cat. 34863), chloroform (cat. C7559), NaCl (cat. S7653), $H_2O_2$ (cat. H1009), SDS (cat. 436143), paraformaldehyde (cat. 158127), glycine (cat. G8898), penicillin G (cat. P7794), streptomycin (cat. S9137), sucrose (cat. S7903), PBS (cat. P5493), xylene (cat. 534056), Tween-20 (cat. P9416), Trizma base (Tris; cat. T1503), polybrene (cat. H9268), ammonium persulfate (APS; cat. A3678), tetramethylethylenediamine (TEMED; cat. T9281), diethylpyrocarbonate (DEPC)-treated water (cat. 693520), ammonium hydroxide solution (cat. 338818), hydrochloric acid in ethanol (cat. 1.00327), trypsin (cat. T1426), Non-fat-Dried Milk bovine (cat. M7409), collagenase B (cat. 11088831001), type I collagenase (cat. C0130), Y-27632 (cat. Y0503), (N-methyl-d3)-palmitoyl-l-carnitine ($d_3$-L-carnitine; cat. 55107), BSA (cat. A2153), N-nitrosodiethylamine (DEN; cat. N0756), Kolliphor HS 15 (cat. 42966), percoll (cat. #GE17-0891-09), Ficoll-Paque Premium (ficoll; cat. GE17-5442-03), LPS (cat. L2630), and corn oil (cat. C8267) were purchased from Sigma. ReverTra Ace qPCR RT Master Mix with gDNA Remover (cat. FSQ-301) was purchased from Toyobo. SignalStain DAB Substrate Kit (cat. 8059) was purchased from Cell Signaling

Technology. LabAssay triglyceride reagent (cat. 290-63701) was purchased from Wako Pure Chemical Industries. Western-Bright ECL and Peroxide solutions (cat. 210414-73) were purchased from Advansta. Acrylamide/Bis Solution (30%), 29:1 (cat. 1610156) was purchased from Bio-Rad. Lipofectamine 2000 (cat. 11668500), GlutaMAX (cat. 35050061), MEM non-essential amino acids solution (cat. 11140050), sodium pyruvate (cat. 11360070), Maxima SYBR Green/ROX qPCR Master Mix (cat. K0223), FBS (cat. 10099141C), William's E medium, no glutamine (cat. 12551032), DMEM-high glucose (cat. 12800082), DMEM/F-12 (cat. 11320033), RPMI 1640 (cat. 11875093), Liver Perfusion Medium (cat. 17701), Liver Digest Medium (cat. 17703), Prestained Protein MW Marker (cat. 26612) and TRIzol (cat. 15596018) were purchased from Thermo Scientific. Cytofix/Cytoperm Fixation/Permeabilization Kit (cat. 554714) was purchased from BD Biosciences. ALT assay kit (cat. C009-2-1) and AST assay kit (cat. C010-2-1) were purchased from Nanjing Jiancheng Bioengineering Institute. Paraplast High Melt Paraffin (cat. 39601095) was purchased from Leica. [U-$^{13}$C]-glutamine (cat. 184161-19-1) was purchased from Cambridge Isotope Laboratories. $CCl_4$ (cat. C805329) was purchased from Macklin. Annexin V Apoptosis Detection Kits (cat. 88-8007-74) was purchased from eBioscience. BsmB I (cat. R0580L), NEBNext Poly(A) mRNA Magnetic Isolation Module (cat. E7490L) and NEBNext Ultra RNA Library Prep Kit for Illumina (cat. E7770L) were purchased from New England Biolabs. Phosphatase Inhibitor Cocktail I (cat. HY-K0021), Phosphatase Inhibitor Cocktail II (cat. HY-K0022), Protease Inhibitor Cocktail (cat. HY-K0010), and DAPI (cat. HY-D0814) were purchased from MedChemExpress. Haematoxylin solution (cat. BSBA-4021A) and eosin Y-solution (cat. ZLI-9613) were purchased from ZSGB-BIO, China. Red Cell Lysis Buffer (cat. RT122) was purchased from Tiangen. TUNEL Apoptosis Detection Kit (cat. 40307ES60) and 0.4% Trypan Blue Solution (cat. 40207ES20) were purchased from Yeason.

**Quantification of chemokine mRNA levels by real-time PCR**

Mice were killed by cervical dislocation, immediately followed by dissection of the tumor and the para-carcinoma tissues. Total RNA was then prepared by lysing the tissue with 1 mL of TRIzol, followed by the addition of 270 μL of chloroform and vigorous mixing. After centrifugation at 12,000*g* for 15 min at 4 °C, 450 μL of the upper aqueous layer was transferred to a clean tube. The RNA was then precipitated by adding 675 μL of isopropanol, followed by centrifugation at 12,000*g* for 15 min at 4 °C. The pellet was washed with 75% ethanol 3 times by centrifugation at 12,000*g* for 5 min, and was dissolved in 200 μL of DEPC-treated water. The concentration of RNA was determined using a NanoDrop 2000 spectrophotometer (Thermo Scientific). A total of 2 μg of RNA was diluted with DEPC-treated water to a final volume of 4 μL at 65 °C for 5 min, and immediately chilled on ice. Random Primer Mix, Enzyme Mix and 5× RT buffer (all from the ReverTra Ace qPCR RT kit) were then added to the RNA solution, followed by incubation at 37 °C for 15 min, and then at 98 °C for 5 min on a thermocycler. The reverse-transcribed cDNA was quantified with Maxima SYBR Green/ROX qPCR master mix on a LightCycler 480 system (Roche) with the following programs: pre-denaturing at 95 °C for 10 min; denaturing at 95 °C for 10 s, then annealing and extending at 60 °C for 30 s in each cycle; cycle number: 40. Primer pairs were designed based on the PrimerBank database (https://pga.mgh.harvard.edu/primerbank/), and their sequences are: mouse *Actb*, 5'-GGCTGTATTCCCCTCCATCG-3' and 5'-CCAGTTGGTAACAATGCCATGT-3'; mouse *Cxcl9*, 5'-TCCTTTTGGGCATCATCTTCC-3' and 5'-TTTGTAGTGGATCGTGCCTCG-3'; mouse *Cxcl10*, 5'-CCAAGTGCTGCCGTCATTTTC-3' and 5'-GGCTCGCAGGGATGATTTCAA-3'; and mouse *Cxcl11*, 5'-GGCTTCCTTATGTTCAAACAGGG-3' and 5'-GCCGTTACTCGGGTAAATTACA-

3'. Data were analyzed using Light-Cycler 96 software (v.1.1, Roche). The mRNA level related to beta-actin (*Actb*) was then calculated using Excel software (2016, Microsoft).

RNA sequencing

Levels of chemokine gene expression in tumor and para-tumor tissues were determined through RNA-sequencing performed by Tissuebank Biotechnology Co., Ltd (Shanghai, China). Briefly, 50 mg of tumor and para-tumor tissue dissected by freeze clamp were instantly lysed in 1 mL of TRIzol, followed by centrifuged at 12,000*g* for 15 min at 4 °C. Some 900 μL of supernatant (without the lipid layer) was transferred to an RNase-free tube, followed by mixing with 270 μL of chloroform. After vigorous vortexing for 15 s, the mixture was centrifuged at 12,000*g* for 15 min at 4°C, and some 450 μL of the upper aqueous layer was transferred to an RNase-free tube. The RNA was then precipitated by adding 675 μL of isopropanol, followed by centrifugation at 12,000*g* for 30 min at 4 °C. The pellet was washed twice with 75% (v/v, in DEPC-treated water) ethanol, and was dissolved with 200 μL of DEPC-treated water. The concentration of RNA was determined by a NanoDrop 2000 spectrophotometer (Thermo), and the integrity of RNA was by the Fragment Analyzer 5400 (Agilent Technologies).

The cDNA libraries were then generated using the NEBNext Poly(A) mRNA Magnetic Isolation Module and the NEBNext Ultra RNA Library Prep Kit for Illumina, following the manufacturer's instructions. Briefly, mRNA from each sample was captured through the NEBNext Oligo $d(T)_{25}$ beads, followed by fragmentation in the First Strand Synthesis Reaction Buffer at 94 °C for 15 min. The fragmented mRNAs were used to synthesize first-strand cDNA with random hexamer primers and M-MuLV Reverse Transcriptase (M-MLV), followed by second-strand cDNA with DNA Polymerase I and RNase H. The remaining overhangs of cDNAs were then converted into blunt ends, followed by adenylating the 3' ends using the NEBNext End Prep Enzyme Mix, and then ligated to the NEBNext Adaptors. The ligation products were then purified with the AMPure XP Beads (Beckman Coulter), during which the cDNA fragments of 250-300 bp in length were enriched, followed by treatment with the USER Enzyme at 37°C for 15 min to remove the hairpin structures on the adaptors. After incubating at 95 °C for 15 min, the adaptor-ligated cDNAs were amplified by PCR using the Phusion High-Fidelity DNA polymerase, the Universal PCR Primers, and the Index Primer, followed by purified with AMPure XP Beads, and the quality of the library was assessed on the Bioanalyzer 2100 (Agilent Technologies). The amplified library was then clustered using the TruSeq PE Cluster Kit v3-cBot-HS kit (Illumia) on a cBot Cluster Generation System (Illumia), followed by sequencing on a Novaseq 6000 platform (Illumia) during which 150-bp paired-end reads were generated.

The original image files generated during sequencing were transformed to short reads (Raw data, in FASTQ format) by base calling. The low-quality sequences (Phred quality score <5), sequences contaminated with adapter (detected in either one read of a paired reads), and the unrecognizable sequences (over 10% bases that were unrecognizable in either one read in a paired reads) were removed from raw reads by the fastp software (version 0.21.1; ref. (*97*)). Expression levels of gene were quantified through FPKM (fragment per kilobase of gene per million reads mapped) values. To acquire the FPKM value of each gene, reads were first mapped to the reference sequence of mice using the Hisat2 software (version 2.2.1) as described previously (*98*) to make sure that the reads could be uniquely mapped to the gene chosen to calculate the FPKM values. For genes with more than one alternative transcript, the longest transcript was selected to calculate the FPKM.

The FPKM was calculated by the feature Counts software (version 2.0.1) as described previously (*99*). FPKM values for *Cxcl9*, *Cxcl10*, and *Cxcl11* genes were plotted using Prism 9 (GraphPad) software.

<u>Determination of aldometanib concentrations in mouse serum and tissues</u>
The concentrations of aldometanib in the liver and serum of DEN-HFD mice were determined using the protocol described previously (*41*), with minor modifications. Briefly, C57BL/6J mice, after intraperitoneal injection with two doses of DEN on postnatal days 28 and 35, and then feeding with HFD starting from day 49, were administered with 100 mg/L aldometanib in the drinking water for 1 week starting after 33 weeks of HFD treatment. At the end of the aldometanib treatment, liver tissues and blood were collected. The blood was left at room temperature for 20 min and then centrifuged at 3,000*g* for 30 min at 4 °C to obtain the serum. Some 100 μL of serum or 100 mg of liver or tumor tissue was lysed with 1 mL of ice-cold 80% (v/v) methanol in water containing 2.3 ng/mL $d_3$-L-carnitine C16:0 as an internal standard, followed by centrifuging at 20,000*g* for 15 min at 4 °C. Some 600 μL of supernatant was collected, lyophilized in a vacuum concentrator (CentriVap Benchtop Centrifugal Vacuum Concentrator, equipped with a CentriVap -84 °C Cold Trap and a Scroll Vacuum Pump, Labconco) at 4 °C, and then dissolved in 100 μL of 70% (v/v, in water) methanol. Samples were analyzed on a QTRAP MS (QTRAP 6500+, SCIEX) interfaced with a UPLC system (Acquity I-class, Waters). Some 2 μL of each sample was loaded onto a reverse-phase column (ACQUITY UPLC BEH C18, 1.7 μm, 2.1 × 50 mm; 186002350, Waters). The mobile phase consisting of 0.1% formic acid in LC-MS-grade water (mobile phase A) and LC-MS-grade methanol (mobile phase B) was run at a flow rate of 0.2 mL/min. The analytes were separated with the following gradient program: 70% B increased to 100% B in 5 min, held for 2 min, and the post time was set to 3 min. The QTRAP mass spectrometer used a Turbo V ion source and ran in positive mode with a spray voltage of 5,500 V, source temperature of 400 °C, gas 1 of 40 psi, gas 2 of 50 psi, and curtain gas of 40 psi. Aldometanib was measured using the multiple reaction monitoring (MRM) mode, and declustering potentials and collision energies were optimized through the use of analytical standards. The following transitions were used for monitoring each compound: 465.3/240.6 and 465.3/159.1 for aldometanib and 403.4/85 for $d_3$-L-carnitine C16:0 as internal standard. Data were collected using Analyst software (v.1.6.3, SCIEX), and the relative amounts of aldometanib were analyzed using MultiQuant software (v.3.0.2, SCIEX).

<u>Immunoblotting</u>
To analyze the levels of p-AMPKα and p-ACC, p-S6K, and the protein levels of cell death markers in cultured cells, cells were grown to 95% confluence in a well of a 6-well dish, and were lysed with 300 μL of ice-cold lysis buffer for each well. To analyze the levels of apoptotic markers in HCC tissues, 100 mg of freshly excised tissue was lysed with ice-cold lysis buffer (10 μL/mg liver weight), followed by homogenization by a hand-hold homogenizer T 10 basic ULTRA-TURRAX equipped with an S 10 N - 5 G dispersing tool, IKA. The lysates were sonicated by a sonicator (VCX130PB, SONICS) equipped with a 5/64" (2 mm) stepped microtip (cat. 630-0423, SONICS) for 3 s at 25% maximum power on ice, and then centrifuged at 20,000*g* for 10 min at 4 °C. After discarding the pellet, an equal volume of 2× SDS sample buffer was added into the supernatant. Samples were then boiled for 10 min before gel electrophoresis and immunoblotting.

All protein samples were subjected to immunoblotting on the same day of preparation without any freeze-thaw cycle.

For IB, the SDS-PAGE gels were prepared in-house as described previously (*35*). Briefly, the resolving gel solution (8%, 10 mL) was prepared by mixing 1.9 mL of 30% Acryl/Bis solution, 1 mL of 10× Lower Buffer (3.5 M Tris, 1% (w/v) SDS, pH 8.8), and 0.48 mL of 65% (w/v) sucrose (dissolved in water) with 6.62 mL of water; and the stacking gel solution (5 mL) was prepared by mixing 668 μL of 30% Acryl/Bis solution, and 1.25 mL of 4× Stacking Buffer (0.5 M Tris, 0.4% (w/v) SDS, pH 6.8) with 3.08 mL water. For each glass gel plate (with 1.0-mm spacer; 1653308 and 1653311, Bio-Rad), approximately 7 mL of resolving gel solution and 2.5 mL of stacking gel solution were required. APS (to 0.1% (w/v) final concentration) and TEMED (to 0.1% (v/v) final concentration) were added to the resolving gel solution. The resolving gel was overlayed with 2 mL of 75% (v/v) ethanol before the acrylamide polymerization. After around 20 min (when a clear line between resolving gel and ethanol is seen), the overlaid ethanol was poured off, dried with filter paper, and placed at room temperature for another 15 min to let the ethanol evaporate completely. The gel cassette was then filled with APS/TEMED-supplemented stacking gel solution, followed by placing a 15-well comb into the cassette, and then placed at room temperature for 20 min. After removing the comb, the gel was rinsed with Running Buffer (25 mM Tris, 192 mM glycine, 1% (w/v) SDS, pH 8.3) before sample loading. Samples of less than 10 μL were loaded into wells, and the electrophoresis was run at 100 V by a Mini-PROTEAN Tetra Electrophoresis Cell (Bio-Rad). All samples were resolved on 8% resolving PAGE gels, except those for caspase-3, caspase-7, GSDME, GSDMD, and PUMA were run on 12% gels (prepared as those of 8%, except that a final concentration of 12% Acryl/Bis was added to the resolving gel solution). The resolved proteins were then electrically transferred to pre-cut PVDF membranes (0.45 μm; IPVH00010, Merck), which were pre-incubated in methanol for 1 min, followed by equilibrating and soaking in pre-cooled Transfer Buffer (25 mM Tris, 192 mM glycine, 10% (v/v) methanol) for more than 5 min. After preparing the gel/membrane sandwich, the transfer was performed at a voltage of 100 V in a Mini Trans-Blot Cell (Bio-Rad) for 1 h at 4 °C. The blotted PVDF membrane was then incubated in blocking buffer (5% (w/v) BSA or 5% (w/v) non-fat milk (according to the instructions from the antibody suppliers) dissolved in TBST ((40 mM Tris, 275 μM NaCl, 0.2% (v/v) Tween-20, pH7.6)) for another 2 h on an orbital shaker at room temperature, followed by rinsing with TBST for twice, 5 min each. The PVDF membrane was incubated with the desired primary antibody overnight at 4 °C on an orbital shaker with gentle shaking, followed by rinsing with TBST three times, 5 min each at room temperature, and then the membrane was incubated with the secondary antibody for 3 h at room temperature with gentle shaking. The secondary antibody was then removed, and the PVDF membrane was further washed with TBST three times, 5 min each at room temperature. PVDF membranes were then incubated in ECL mixture (by mixing equal volumes of ECL solution and peroxide solution for 5 min), then were placed in plastic wrap and laid with Medical X-Ray Film (FUJIFILM) in a light-proof cassette for a desired period of time. The films were then developed with X-OMAT MX Developer and Replenisher and X-OMAT MX Fixer and Replenisher solutions (Carestream) on a Medical X-Ray Processor (Carestream) using Developer (Model 002, Carestream). The developed films were scanned using a Perfection V850 Pro scanner (Epson) using Epson Scan software (v.3.9.3.4, Epson), and were cropped using Photoshop (2023, Adobe). Levels of total proteins and phosphorylated proteins were analyzed on separate gels, and representative immunoblots are shown. The band intensities on developed films were quantified using ImageJ (v.1.8.0, National Institutes of Health Freeware). Uncropped immunoblots are uploaded as a “Full scans” file.

Measurement of adenylates

To analyze ATP, ADP and AMP in tissues, HPLC-MS was performed as described previously (*41*). In brief, 100 mg of liver tissue dissected by freeze clamp were instantly lysed in 1 mL of methanol, then mixed with 1 mL of chloroform and 400 μL of water (containing 4 μg/mL [U-$^{13}$C]-glutamine), followed by 20 s of vortexing. After centrifugation at 15,000*g* for another 15 min at 4 °C, 800 μL of aqueous phase was collected, lyophilized in a vacuum concentrator at 4 °C, and then dissolved in 30 μL of 50% (v/v, in water) acetonitrile. Measurement of AMP and ATP level was based on ref. (*100*) using a QTRAP MS (SCIEX, QTRAP 5500) interfaced with a UPLC system (SCIEX, ExionLC AD). A total of 2 μL of each sample was loaded onto a HILIC column (ZIC-pHILIC, 5 μm, 2.1 × 100 mm, PN: 1.50462.0001, Millipore). The mobile phase consisted of 15 mM ammonium acetate containing 3 mL/L ammonium hydroxide (>28%, v/v) in the LC-MS-grade water (mobile phase A) and LC-MS-grade, 90% (v/v) acetonitrile in LC-MS-grade water (mobile phase B) run at a flow rate of 0.2 mL/min. AMP, ADP, and ATP were separated with the following HPLC gradient elution program: 95% B held for 2 min, then to 45% B in 13 min, held for 3 min, and then back to 95% B for 4 min. The mass spectrometer was run on a Turbo V ion source in negative mode with a spray voltage of -4,500 V, source temperature at 550 °C, gas no.1 at 50 psi, gas no.2 at 55 psi, and curtain gas at 40 psi. The following transitions were used for monitoring each compound: 505.9/158.9 and 505.9/408.0 for ATP; 425.9/133.9, 425.9/158.8, and 425.9/328.0 for ADP; 345.9/79.9, 345.9/96.9 and 345.9/133.9 for AMP; and 149.9/114 for [U-$^{13}$C]-glutamine. Data were collected using Analyst 1.7.1 software (SCIEX), and the relative amounts of metabolites were analyzed using MultiQuant 3.0.3 software (SCIEX). Note that a portion of ADP and ATP could lose one or two phosphate groups during in-source fragmentation, thus leaving the same m/z ratios as AMP and ADP, which were corrected according to their different retention times in the column.

Determination of the composition of immune cells in HCC tissues

The composition of immune cells in HCC tissues was determined by a flow-cytometry-based method as described previously (*101, 102*), with minor modifications. Briefly, the HCC-bearing mice were euthanized by cervical dislocation, and the HCC tissues were quickly excised without draining the blood. The tissues were then minced into 1-mm pieces using ophthalmic scissors, and approximately 100 mg of tissues were digested with 2 mL of type I collagenase (1 mg/mL) dissolved in PBS supplemented with 10% FBS in a 5-mL conical tube for 60 min at 37 ℃ in a shaker at 50 rpm. The digestions were then filtered using a 70-μm cell strainer, followed by centrifuge of the filtrate at 400*g* for 5 min at 25 ℃. The yielded cell pellets were gently mixed in 5 mL of 40% percoll (prepared by mixing 2 mL of percoll with 0.22 mL of 10× PBS and 2.78 mL of 1× PBS), and then overlayered on top of a 3 mL of 80% percoll cushion (consisting of 2.4 mL percoll, 0.27 mL of 10× PBS, and 0.33 mL 1× PBS) in a 15-mL conical tube, followed by centrifuge for 20 min at 2000*g* at 25 ℃. Note that the centrifuge should be set at minimum acceleration and deactivate the brake program, as it may disturb the percoll gradient and, in turn, lower the cell yield. After centrifugation, the fraction located at the 40%-80% percoll interface, approximately 2 mL, which contained immune cells, was collected by a Pasteur pipet and mixed with 10 mL of PBS solution, and then centrifuged for 5 min at 400*g* at 25 ℃. Some 1 × 10$^6$ cells were then incubated with the anti-CD16/CD32 antibody (1:100 in PBS, in a total volume of 50 μL), for 30 min at room temperature in a 96-well round bottom plate (non-tissue culture treated, cat. CLS7007-24EA, Corning). The desired combinations of primary antibodies, in a total volume of 50 μL as specified in the figure legends, were then added to the cells and incubated for another 30 min at room temperature, followed by washing with 200 μL of PBS twice. In particular, before staining granzyme B, IFNγ, IL-1β, IL-4, and IL-17a, cells were treated with 200 μL of fixation and permeabilization buffer included in the Cytofix/Cytoperm Fixation/Permeabilization Kit for 30 min at room temperature, followed by one wash with 200 μL of washing buffer. Cells were incubated in 200 μL of 1% (v/v) formalin in the dark at 4 °C before the flow cytometry analysis.

Flow cytometry was conducted using an LSRFortessa X20 cell analyzer (BD Biosciences), which is equipped with 5 solid-state lasers (355 nm, 15 mW; 405 nm, 50 mW; 488 nm, 50 mW; 561 nm, 30 mW; and 640 nm, 40 mW), a forward scatter (FSC) detector, a side scatter (SSC) detector, and an 18-channel fluorophore detector. In this study, the 488-nm laser and the 530/30 filter were utilized to excite and detect the fluorescence of Alexa Fluor 488, the 488-nm laser and the 710/50 filter for PerCP-Cy5.5, the 561-nm laser and the 586/15 filter for PE, and the 640-nm laser and the 670/14 filter for APC. Detector voltages were optimized using a modified voltage titration approach (*103*). Gating strategies used for identifying each type of immune cells during the analysis were shown in the corresponding panels. Gate boundaries were set either based on control samples or followed density distributions based on best practices. Data were collected by the FACSDiva software (v8.0.2, BD Biosciences), followed by exporting in the FCS 3.1 format. The numbers of each type of immune cell were quantified with the FlowJo software (v10.4.0, BD Biosciences). During the analysis, a combination of manual gating and computational analysis approaches (*104*) was used.

Histology

For H&E staining, liver tissues excised from blood-drained mice were cut into pieces, and were fixed in 4% (v/v) paraformaldehyde within 48 h at room temperature, then transferred to embedding cassettes. The cassettes were then washed in running water for 12 h, followed by successive soaking each for 1 h in 70% ethanol (v/v in water), 80% ethanol, and 95% ethanol. The fixed tissues were further dehydrated in anhydrous ethanol for 1 h twice, followed by immersing in 50% xylene (v/v in ethanol) for 30 min, two changes of xylene, 15 min each; and two changes of paraffin wax (58-60 °C), 1 h each. The dehydrated tissues were embedded in paraffin on a HistoCore Arcadia Paraffin Embedding Machine (Leica). Paraffin blocks were then sectioned at a thickness of 4 μm, dried on an adhesion microscope slide, followed by rehydrating in the following order: two changes of xylene at 70 °C, 10 min each; two changes of anhydrous ethanol, 5 min each; two changes of 95% ethanol, 5 min each; one change each for 5 min of 80% ethanol, 70% ethanol, and 50% ethanol, and then briefly in water. The sections were then stained in hematoxylin solution for 8 min, then washed in running water for 5 min, differentiated in 1% hydrochloric acid (in ethanol) for 30 s, washed in running water for 1 min, and immersed in 0.2% (v/v in water) ammonium hydroxide solution for 30 s, washed in running water for 1 min, and stained in eosin Y-solution for 30 s. The stained sections were dehydrated in 70% ethanol for 5 min; twice in 95% ethanol, 5 min each; twice in anhydrous ethanol, 5 min each; and two changes of xylene, 15 min each. The stained sections were mounted with Canada balsam and visualized using a Zeiss Observer Z1 (Carl Zeiss, Jena, Germany).

For immunohistochemistry (IHC), liver tissues were fixed, dehydrated, embedded, sectioned, and re-hydrated as in H&E staining and were washed with water three times, 5 min each, at room temperature. The sections were then incubated in pre-heated (approximately 95 °C) citrate antigen retrieval buffer (1 mM sodium citrate, pH 6.0, 0.05% (v/v) Tween-20) for 1.5 min, followed by cooling at room temperature for another 60 min. The sections were then washed with washing buffer (0.1% (v/v) Tween-20 in PBS) twice, 5 min each at room temperature, and then incubated in 10% (w/w) $H_2O_2$ (in methanol) solution at room temperature for 5 min, followed by washing with washing buffer for three times, 5 min each at room temperature. The sections were then incubated in 1% (w/v) BSA (diluted with PBS), at room temperature for 20 min. After draining, the liver sections were circled by a PAP pen (cat. #Z377821, Sigma), followed by incubation with primary antibodies (diluted in 1% BSA solution) at 4 °C in a dark, humidified chamber, followed by washing with washing buffer for 3 times, 5 min each at room temperature. The sections were then incubated with HRP-conjugated goat anti-rabbit IgG or goat anti-mouse IgG (diluted in 1% BSA solution) for 1h at room temperature in a dark, humidified chamber, followed by washing with washing buffer for 3 times, 5 min each at room temperature. The sections were then incubated with the DAB working solution, which was freshly prepared by adding 30 μL of SignalStain DAB Chromogen Concentrate to 1 mL of SignalStain DAB Diluent (both included in the SignalStain DAB Substrate Kit), and then mixed well. The incubation lasted for approximately 5 min in the dark at room temperature, until a good staining intensity developed, as checked on a Zeiss Observer Z1 (Carl Zeiss, Jena, Germany). The sections were then washed with running water for 5 min, followed by staining in hematoxylin solution for 8 min, and then washed in running water for another 5 min. They were then differentiated in 1% hydrochloric acid (in ethanol) for 30 s, washed in running water for 1 min, and dehydrated in 70% ethanol for 5 min, followed by two washes in 95% ethanol for 5 min each, two washes in anhydrous ethanol for 5 min each, and finally two changes of xylene, each for 15 min. The stained sections were mounted with Canada balsam and visualized using an Axioscan 7 microscope (Zeiss).

For measuring hepatic TAG contents, mice were euthanized by cervical dislocation, and the livers were immediately excised and rinsed in PBS three times. Some 50 mg of tissue was homogenized with a hand-held homogenizer (T 10 basic ULTRA-TURRAX equipped with an S10N-5G dispersing tool, IKA) in 1 mL of PBS containing 5% (v/v) Triton X-100 and sonicated by a sonicator (VCX130PB equipped with a 5/64" stepped microtip, SONICS) at 30% maximum power for 20 cycles, 1 s per cycle with 2 s intermittent at room temperature. Some 600 μL of the supernatant was centrifuged at 20,000*g* for 10 min at 25 °C, and the supernatant was boiled for 20 min, followed by centrifugation at 20,000*g* at 25 °C for 10 min. Some 3 μL of the supernatant was used for each test, using the LabAssay triglyceride kit according to the manufacturer's instructions.

Determination of serum ALT and AST levels

Serum samples were freshly prepared as described in the "HCC-HFD, *MYC*;*Trp53*$^{-/-}$ mice and HCC xenografts" section. Levels of serum ALT and AST were determined using the ALT assay kit and AST assay kit following the manufacturer's instructions. Some 5 μL of serum was used for each test, and the optical density (OD) was recorded by a SpectraMax M5 microplate reader (Molecular Devices) using the SoftMax Pro software (v.5.4.1.1, Molecular Devices).

Determination of the rates of cell death

Effects of aldometanib on cell death in cultured cells were determined by a flow cytometry-based method, as described previously (*79*), with minor modifications. Briefly, HCC cells and or normal liver cells grown to 70-80% confluence in a 6-well dish were treated with aldometanib, followed by washing with 1 mL of PBS (pre-heated to 37 °C), and then trypsinized with 0.25% trypsin, followed by centrifugation at 1,200*g* for 5 min at 4 °C. Some $1 \times 10^6$ trypsinized cells were washed with 0.2 mL of Binding Buffer provided in the Annexin V Apoptosis Detection Kit. Cell suspensions were centrifuged at 1,200*g* for 5 min at 4 °C, and then resuspended with 200 μL of Binding Buffer, followed by staining with 5 μL of Annexin V-APC solution for another 30 min at room temperature in the dark. Cell suspensions were then incubated with 5 μL of PI solution for another 5 min at room temperature in the dark, followed by centrifugation at 1,200*g* for 5 min at 4 °C to remove dyes that are not bound to cells. Cell suspensions were then diluted with 400 μL of Binding Buffer, and then immediately subjected to flow cytometry analysis. Flow cytometry was performed on a BD LSRFortessa X20 cell analyzer, with the 640-nm (40 mW) laser and the 670/14 filter used to excite and detect the fluorescence of Annexin V-APC, and the 561-nm (30 mW) laser and 585/42 filter PI. Detector voltages were optimized using a modified voltage titration approach (*103*). Gating strategies used during the analysis were: a) the FSC-H and SSC-H for selecting intact cells, and b) FSC-H and FSC-A for excluding doublets. Gate boundaries were set either based on control samples, or followed density distributions based on best practices. Data were collected by the FACSDiva software (v8.0.2, BD Biosciences), followed by exporting in the FCS 3.1 format. The numbers of early apoptotic cells (Annexin V positive and PI negative populations), and late apoptotic and necroptotic cells (Annexin V positive and PI positive populations) were quantified with the FlowJo software (v10.4.0, BD Biosciences). For this experiment, a combination of manual gating and computational analysis approaches (*104*) was used.

Effects of aldometanib on cell death in HCC tissues were determined by the TUNEL assay using the TUNEL Apoptosis Detection Kit according to the manufacturer's instructions, with minor modifications. Briefly, the liver paraffin sections were baked at 70 °C for 4 h, followed by quickly immersed with two changes of xylene at room temperature, each for 5 min, and then washed in anhydrous ethanol twice at room temperature, each for 5 min. The sections were then sequentially incubated in 90%, 80%, and 70% ethanol, each for 3 min, followed by briefly washing with PBS. After draining, the sections were circled by a PAP pen, followed by incubation with 100 μL of proteinase K solution (20 μg/mL final concentration, in PBS) at room temperature for 20 min, and then rinsed with PBS for another 5 min at room temperature. The sections were incubated with 100 μL of 1× Equilibration Buffer (freshly prepared by diluting 5× Equilibration Buffer with didistilled water) at room temperature for 30 min, followed by incubation with 100 μL of TdT Reaction Buffer (freshly prepared by mixing didistilled water, 5× Equilibration Buffer, FITC-12-dUTP Labeling Mix, and Recombinant TdT Enzyme at a ratio of 34:10:5:1) in the dark, humidified chamber at 37 °C for 1 h. The sections were then rinsed with PBS for 5 min at room temperature, followed by incubating in 100 μL of 0.1% (v/v) Triton X-100 solution (in PBS containing 5 mg/mL BSA) three times, each for 5 min at room temperature, and then rinsed with PBS for 5 min at room temperature. The sections were then incubated with freshly prepared DAPI solution (2 μg/mL, in PBS) for 5 min in the dark at room temperature, followed by rinsing with didistilled water three times, each for 5 min at room temperature. The drained sections were mounted with Canada balsam and visualized using an Axioscan 7 microscope (Zeiss).

Effects of aldometanib on cell growth were determined by quantifying the numbers of viable cells at different time points during culture using the Trypan Blue staining. Specifically, $2 \times 10^5$ BNL cells, $1.5 \times 10^5$ Huh7 cells, $2 \times 10^5$ JHH-7 cells, $1 \times 10^6$ mouse primary HCC cells, and $4 \times 10^5$ Hepa1-6 cells were seeded into 6-well dishes. After 12 h of incubation, cells were treated with aldometanib at desired concentrations for another 24 h, followed by rinsing with PBS and then trypsinization. The trypsinized cells were collected by centrifuge at 2000*g*, 5 min at room temperature, followed by resuspension with 500 μL of PBS. Some 100 μL of cell suspension was incubated with 100 μL of 0.4% Trypan Blue Solution for 10 min at room temperature, followed by the addition of 20 μL of stained cells to the Millicell Hemocytometer (cat. MDH-2N1, Sigma) to quantify the number of living cells.

<u>Statistical analysis</u>

Statistical analyses were performed using Prism 9 (GraphPad Software), except for the survival curves, which were analyzed using SPSS 27.0 (IBM) by log-rank (Mantel-Cox) test. Each group of data was subjected to the Kolmogorov-Smirnov test, Anderson-Darling test, D'Agostino-Pearson omnibus test, or Shapiro-Wilk test for normal distribution when applicable. An unpaired two-sided Student's *t*-test was used to determine the significance between two groups of normally distributed data. Welch's correction was used for groups with unequal variances. An unpaired two-sided Mann-Whitney test was used to determine the significance between data without a normal distribution. For comparison between multiple groups with one fixed factor, an ordinary one-way ANOVA was used, followed by Tukey's or Dunnett's multiple comparisons test as specified in the legends. For comparison between multiple groups with two fixed factors, an ordinary two-way ANOVA was used, followed by Tukey's or Sidak's multiple comparisons test as specified in the legends. The assumptions of homogeneity of error variances were tested using F-test ($P > 0.05$). The adjusted means and s.e.m. were recorded when the analysis met the above standards. Differences were considered significant when $P < 0.05$, or $P > 0.05$ with large differences of observed effects (as suggested in refs. (*105, 106*)).

### Supplementary Figures

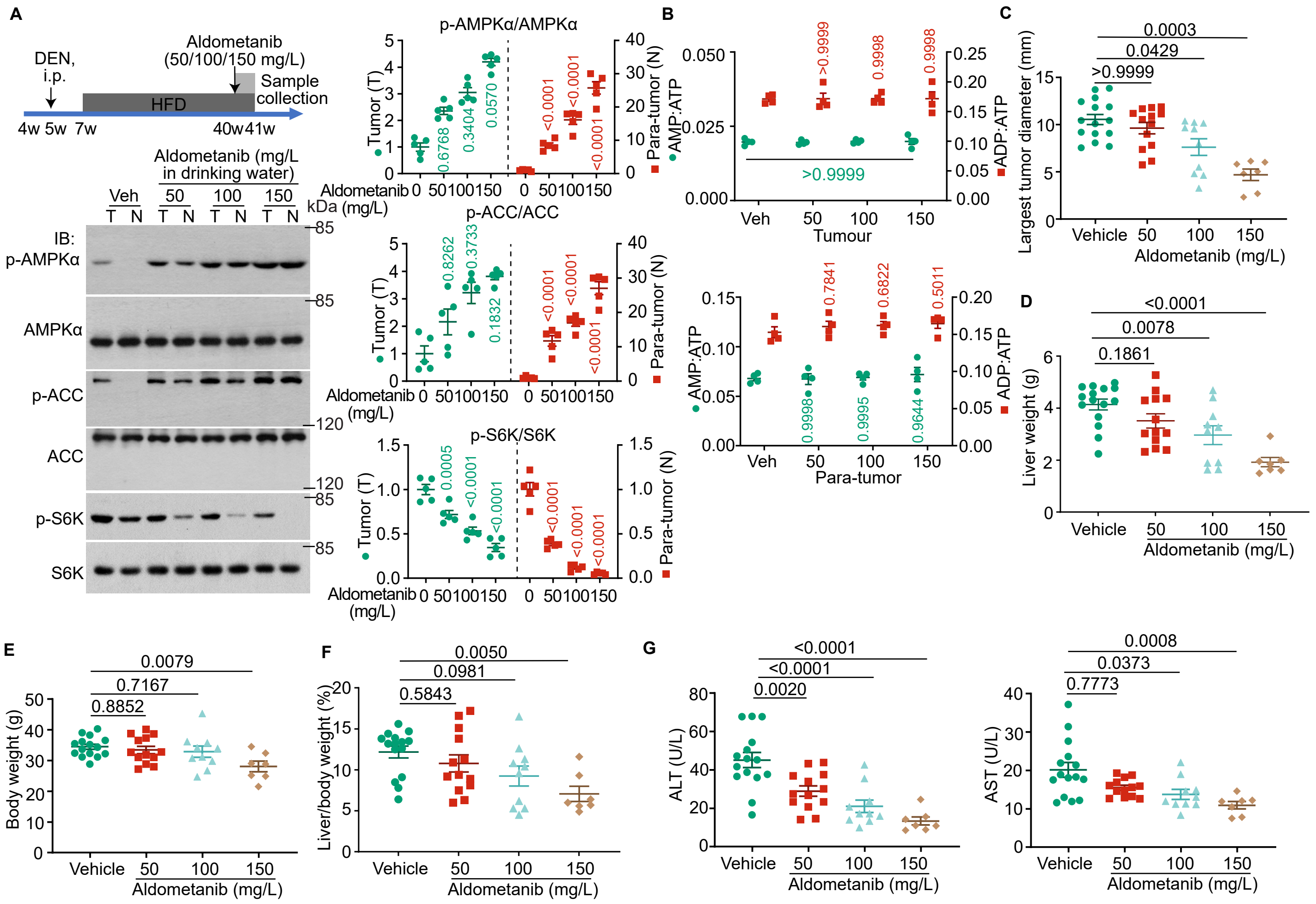


**Extended Data Fig. 1 Aldometanib inhibits hepatocellular carcinomas in a dose-dependent manner.**
**a**, **b**, Aldometanib activates AMPK in the liver tissues of DEN-HFD mice. Wildtype C57BL/6 mice (4 weeks old) were intraperitoneally injected with DEN once a week for 2 weeks, followed by feeding with HFD 2 weeks later (depicted in the upper left panel of **a**). At 40 weeks old (after the formation of HCC, validated in Extended Data Fig. 2a), mice were treated with aldometanib dissolved in drinking water at 50, 100 or 150 mg/l for 7 days. Mice were then euthanised, and the HCC (tumour; T) and para-HCC (para-tumour; N) tissues were freeze-clamped. AMPK activation and mTORC1 inhibition were determined by immunoblotting (**a**, representative blots are shown on the left; the band intensities of blots from five independent experiments were quantified to calculate the ratios of p-AMPKα/AMPKα, p-ACC/ACC and p-S6K/S6K, and are shown on the right panel (means ± s.e.m., $n = 5$ mice for each treatment, with $P$ values calculated by two-way analysis of variance (ANOVA), followed by Tukey's test)), and the ratios of AMP:ATP and ADP:ATP were determined by HPLC-MS (**b**, data are means ± s.e.m., $n = 4$ mice, and $P$ values calculated by two-way ANOVA, followed by Tukey's test).
**c**-**g**, Mice were induced to develop HCC using DEN and HFD, and treated with aldometanib as in Fig. 1b, followed by collection of tissue (**c**-**f**) and serum (**g**) samples at week 48 of age. The largest tumour diameters (**c**), liver weights (**d**), body weights (**e**), liver:body weight ratios (**f**), and serum ALT (**g**, left panel) and AST (**g**, right panel) were then determined. Data are shown as means ± s.e.m., $n = 15$ (vehicle), 13 (50 mg/l), 10 (100 mg/l), or 7 (150 mg/l) mice, with $P$ values calculated by one-way ANOVA, followed by Dunnett's test (**d**, **e**, **f**, and left panel of **g**), or by Kruskal-Wallis test, followed by Dunn's test (**c**, and right panel of **g**).
Experiments in this figure were performed three times.

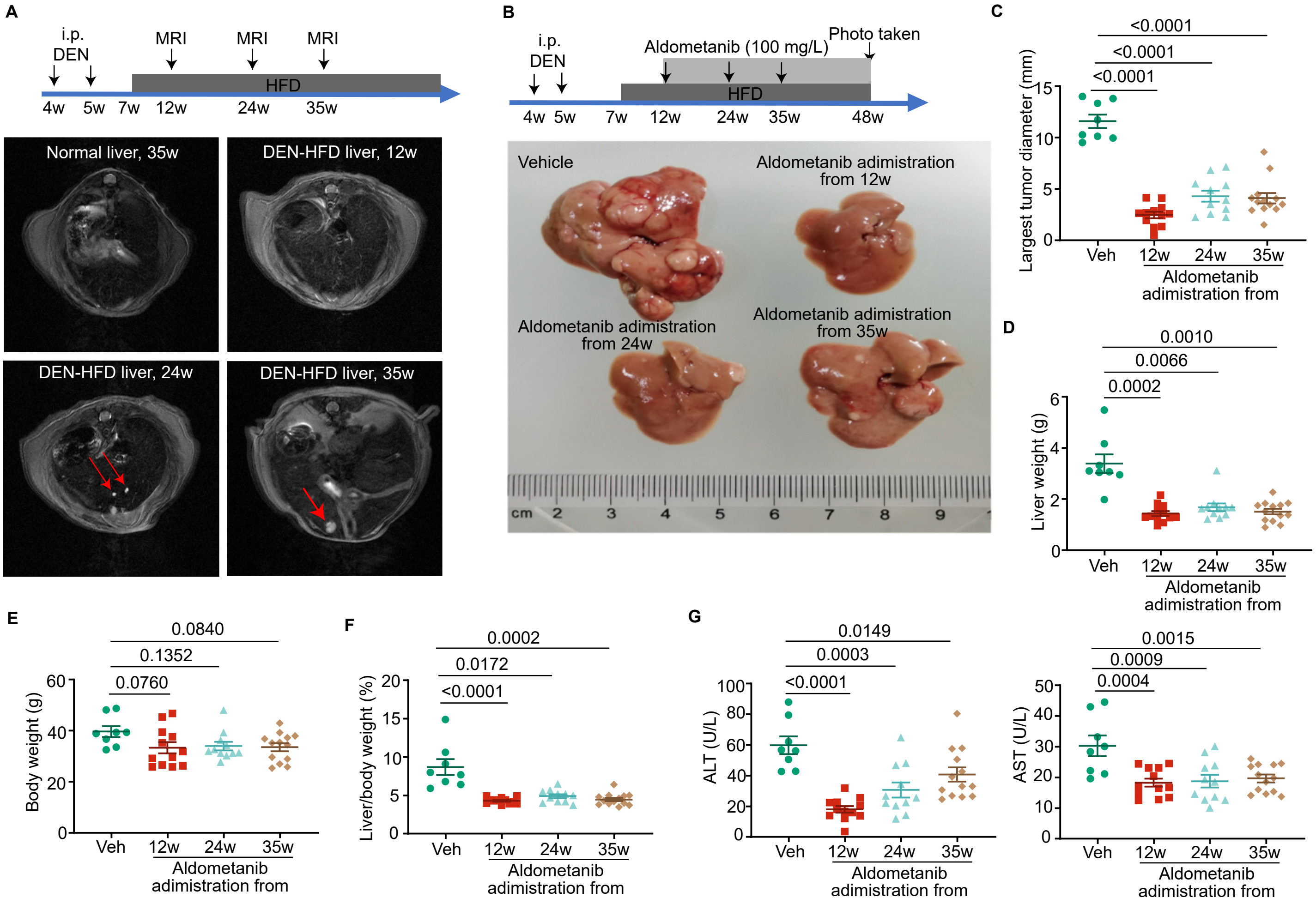


**Fig. S2 Aldometanib is effective in inhibiting late-stage HCC.**
(**A**) Timeline for the development of HCC in the DEN-HFD mice. Mice were induced to develop HCC using DEN and HFD as in Fig. 1A. At 12, 24, and 35 weeks old, the mice were subjected to MRI (depicted in the upper panel, using the DEN- and HFD-untreated normal mice as controls). Representative MRI images are shown in the lower panel, with red arrows pointing to solid HCC tumors.
(**B** to **G**) Aldometanib effectively inhibits late-stage HCC. The DEN-HFD mice were treated with aldometanib as in Fig. 1C. The appearance of liver tissues (B), largest tumor diameters (C), liver weights (D), body weights (E), liver:body weight ratios (F), and serum ALT (G, left panel) and AST (G, right panel) were determined. Data are shown as means ± s.e.m., $n$ = 8 (vehicle), 12 (12 weeks old), 11 (24 weeks old), or 13 (35 weeks old) mice, with $P$ values calculated by one-way ANOVA, followed by Dunnett's test ((C), (E), and (G)), or by Kruskal-Wallis test, followed by Dunn's test ((D) and (F)).
Experiments in this figure were performed three times.

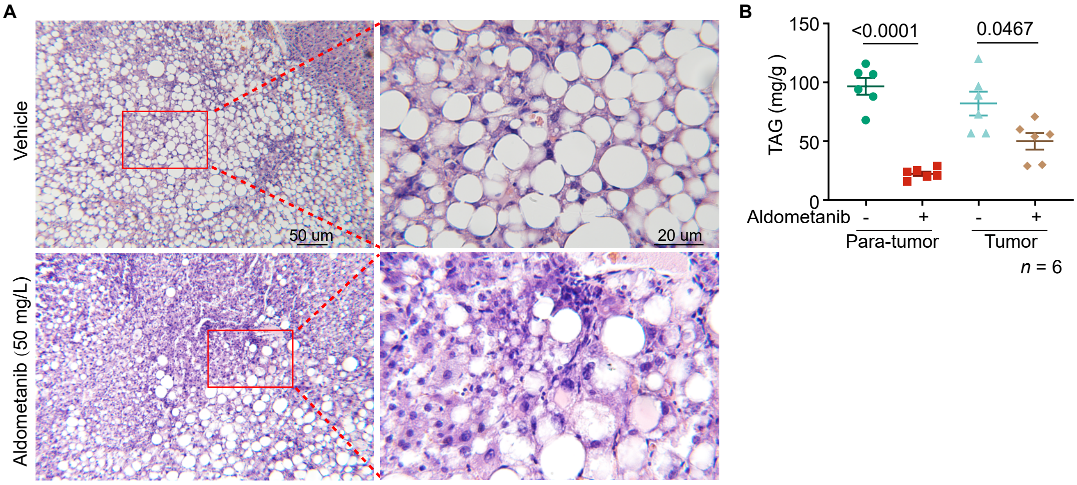


**Fig. S3 Aldometanib decreases triglyceride levels in the liver of DEN-HFD mice.**

(**A** and **B**) The DEN-HFD mice were treated with aldometanib as in Fig. 1B, followed by the collection of liver tissue samples at week 48 of age. Representative images from H&E staining (A) and the triglyceride (TAG) content ((B); data are shown as means ± s.e.m., $n$ = 6 mice, with $P$ values calculated by two-way ANOVA, followed by Tukey) of the liver tissues are shown.

Experiments in this figure were performed three times.

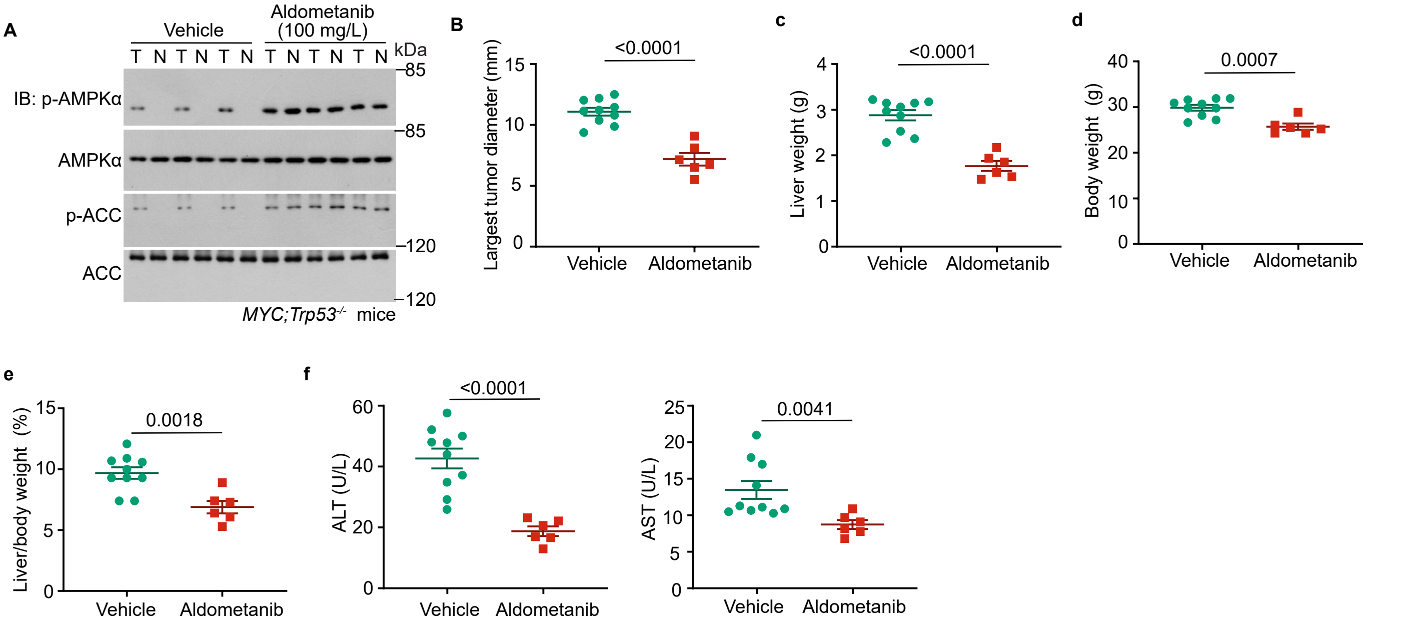


**Fig. S4 Aldometanib inhibits HCC in *MYC*;*Trp53*$^{-/-}$ mice.**

(**A** to **F**) The *MYC*;*Trp53*$^{-/-}$ HCC mice were established and treated with aldometanib as in Fig. 1E. At week 18, mice were euthanized, followed by determination of AMPK activation (A), largest tumor diameters (B), liver weights (C), body weights (D), liver:body weight ratios (E), and serum ALT (F, left panel) and AST (F, right panel). Data are shown as means ± s.e.m., $n$ = 10 (vehicle) or 6 mice (aldometanib), with $P$ values calculated by two-sided Student's $t$-test ((B) to (E)), or by two-sided Student's $t$-test with Welch's correction (F). Experiments in this figure were performed three times.

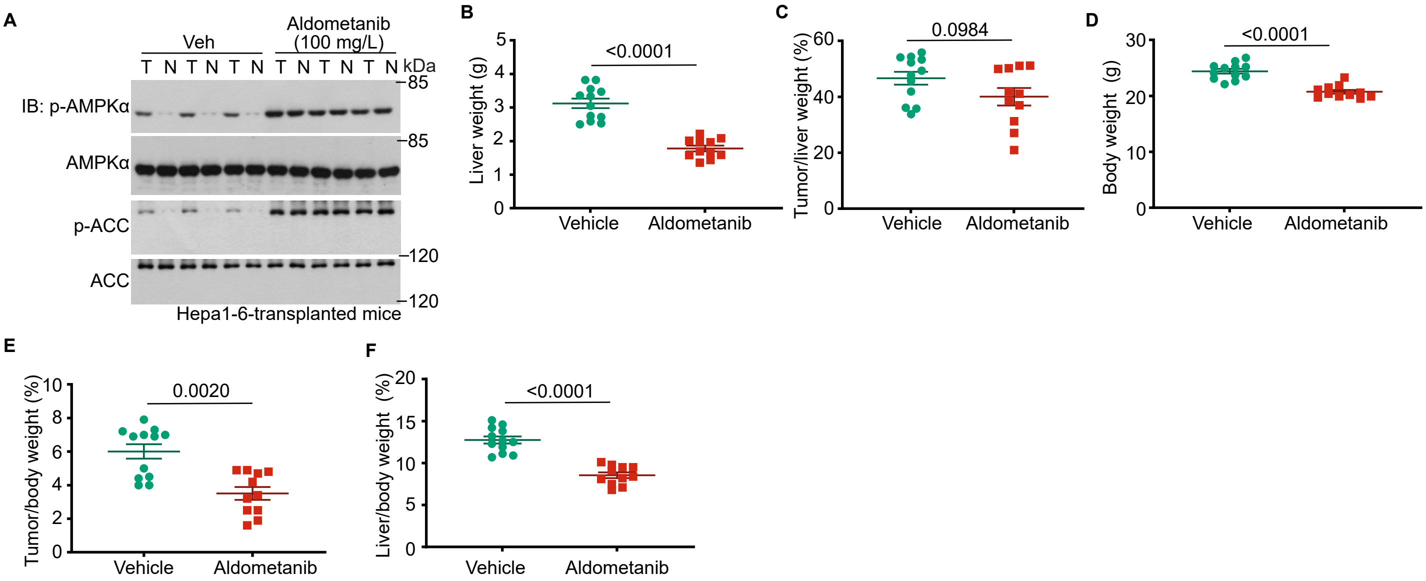


**Fig. S5 Aldometanib inhibits HCC xenografts.**

(**A** to **F**) Mice were transplanted with Hepa1-6 cells into the left liver lobes to develop solid tumors, followed by treatment with aldometanib, as in Fig. 1F. At day 17, mice were euthanized, followed by determination of AMPK activation (A), liver weights (B), tumor:liver weight ratios (C), body weights (D), tumor:body weight ratios (E), and liver:body weight ratios (F). Data are shown as means ± s.e.m., $n$ = 12 (vehicle) or 11 mice (aldometanib), with $P$ values calculated by two-sided Student's $t$-test ((B), (C), (D), and (F)), or by two-tailed Mann–Whitney test (E). Experiments in this figure were performed three times.

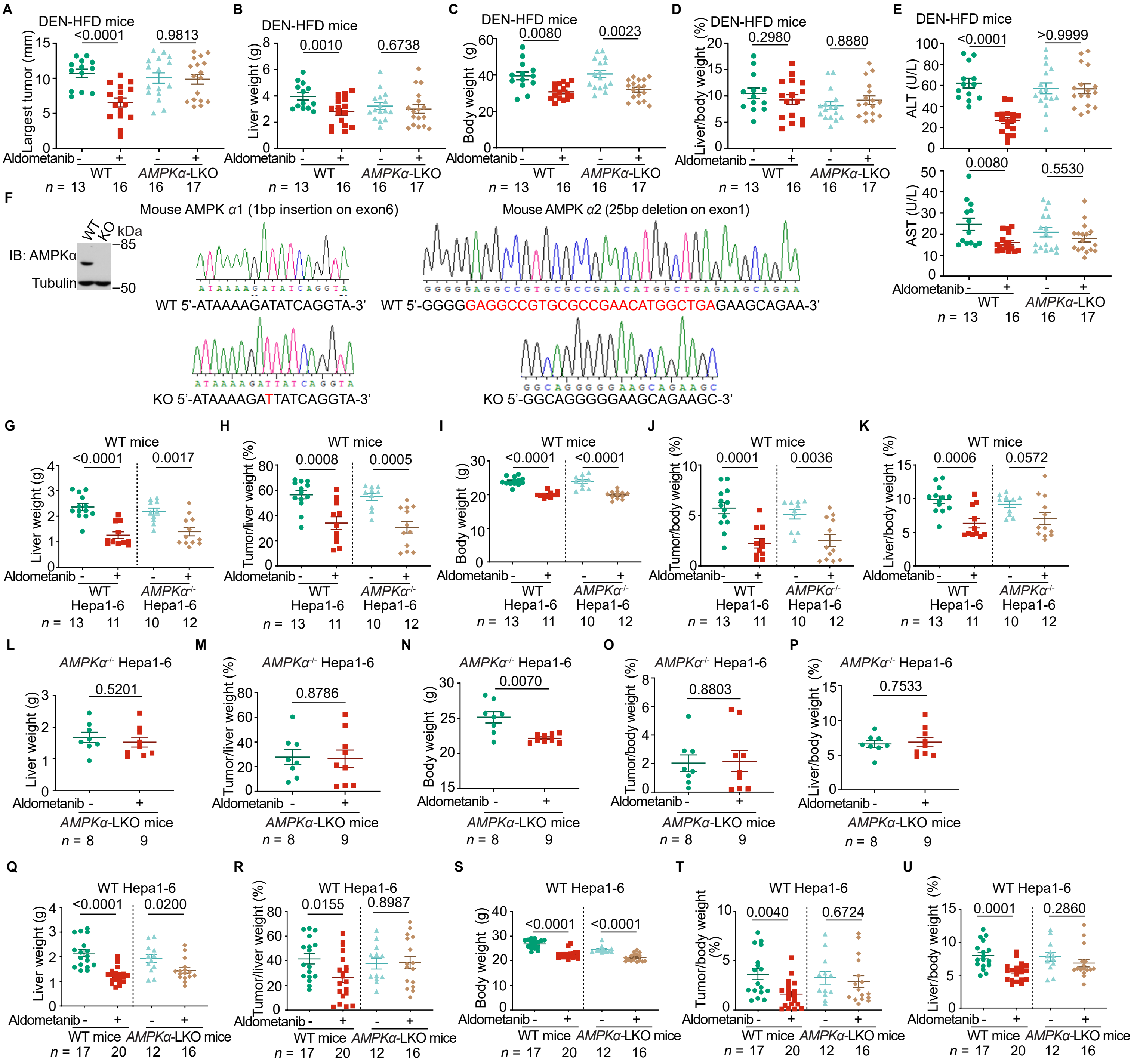


**Fig. S6 AMPK in para-HCC plays a dominant role in aldometanib-induced suppression of HCC.**

(**A to E**) Liver-specific knockout of AMPKα impairs aldometanib-induced suppression of HCC in DEN-HFD mice. Mice were induced to develop HCC using DEN and HFD and were treated with aldometanib as in Fig. 1A, followed by determination of the largest tumor diameters (A), liver weights (B), body weights (C), liver:body weight ratios (D), and serum ALT (E, left panel) and AST (E, right panel). Data are shown as means ± s.e.m., *n* represents the number of mice, and are labeled in each panel, with *P* values calculated by two-way ANOVA, followed by Tukey.

(**F to K**) Aldometanib can still inhibit HCC xenografts of *AMPKα*$^{-/-}$ Hepa1-6 cells. Mice were transplanted with Hepa1-6 cells and treated with aldometanib as in Fig. 2D, followed by determination of liver weights (G), tumor:liver weight ratios (H), body weights (I), tumor:body weight ratios (J), and liver:body weight ratios (K). Data are shown as means ± s.e.m., *n* represents the number of mice, and are labeled in each panel, with *P* values calculated by two-sided Student's *t*-test ((G), (H), (I), (J), and left panel of (K)), or by two-sided Student's *t*-test with Welch's correction (right panel of (K)). See also the validation data for the *AMPKα*$^{-/-}$ Hepa1-6 cells in (F), as determined by immunoblotting (left panel) and sequencing (right panel).

(**L to P**) Aldometanib fails to inhibit HCC xenografts of *AMPKα*$^{-/-}$ Hepa1-6 cells transplanted into the *AMPKα*-KO liver. Mice were transplanted with Hepa1-6 cells and treated with aldometanib as in Fig. 2F, followed by determination of the liver weights (L), tumor: liver weight ratios (M), body weights (N), tumor:body weight ratios (O), and liver:body weight ratios (P). Data are shown as means ± s.e.m., *n* = 8 (vehicle) or 9 (aldometanib) mice, with *P* values calculated by two-sided Student's *t*-test ((L), (M), (O), and (P)), or by two-sided Student's *t*-test with Welch's correction (N).

(**Q** to **U**) Aldometanib fails to inhibit HCC xenografts of wildtype Hepa1-6 cells transplanted into the *AMPKα*-KO liver. Mice were transplanted with Hepa1-6 cells and treated with aldometanib as in Fig. 2H, followed by determination of the liver weights (Q), tumor:liver weight ratios (R), body weights (S), tumor:body weight ratios (T), and liver:body weight ratios (U). Data are shown as means ± s.e.m., *n* represents the number of mice, and are labeled in each panel, with *P* values calculated by two-sided Student's *t*-test (right panels of (Q) and (T), and (R), (S), (U)), or by two-sided Student's *t*-test with Welch's correction (left panel of (Q), and left panel of (T)).

Experiments in this figure were performed three times.

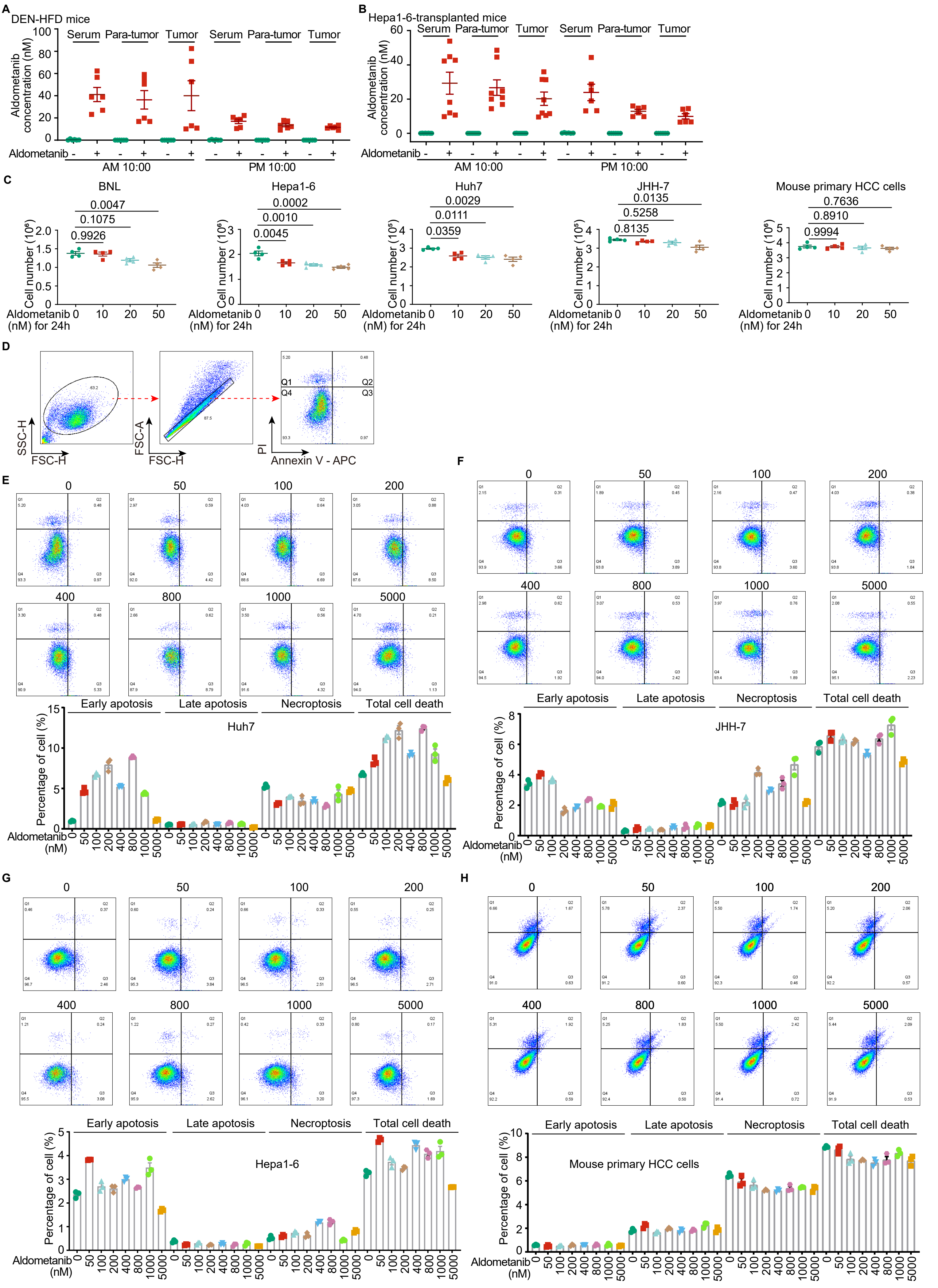
A
DEN-HFD mice
Serum
Para-tumor
Tumor
Aldometanib concentration (nM)
Aldometanib
AM 10:00
PM 10:00
B
Hepa1-6-transplanted mice
C
BNL
Hepa1-6
Huh7
JHH-7
Mouse primary HCC cells
Cell number (10⁶)
Aldometanib (nM) for 24h
D
SSC-H
FSC-H
FSC-A
PI
Annexin V - APC
E
F
Early apoptosis
Late apoptosis
Necroptosis
Total cell death
Percentage of cell (%)
Aldometanib (nM)
G
H

Fig. S7 (cont.)

I

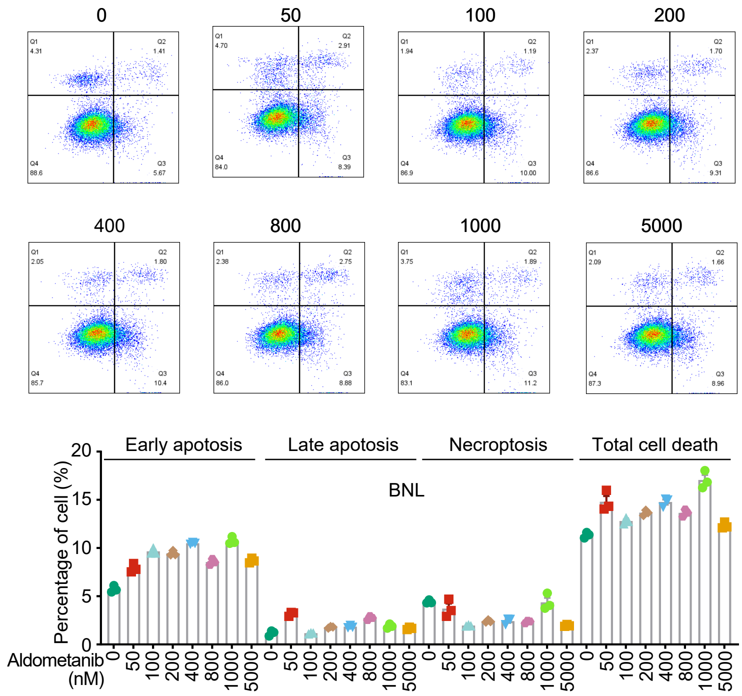


**Fig. S7 Aldometanib exhibits little cytotoxicity to cells in culture.**

(**A** and **B**) Detected concentrations of aldometanib in the serum, liver, and HCC tissues of aldometanib-treated DEN-HFD mice (A) and Hepa1-6-transplanted mice (B). Mice were induced to develop HCC using DEN and HFD, as in Fig. 1A (A), or transplanted with Hepa1-6 cells into the left lobe of the liver, as in Fig. 1F (B). Mice were then treated with 100 mg/L aldometanib at 40 weeks old. At 41 weeks old (A), or 1 week after the administration of aldometanib (B), mice were euthanized, followed by the determination of aldometanib concentrations in the serum, liver, and HCC tissues (tumor) at the indicated time points. Data are shown as means ± s.e.m., $n$ = 8 (10:00 a.m. of B) or 6 (others) mice.

(**C**) Aldometanib slightly inhibits the growth of HCC cells. The normal mouse liver cell line BNL, HCC cell lines Hepa1-6, Huh7, and JHH-7 cells, as well as the primary HCC cells derived from mouse HCC tissues, were treated with aldometanib at indicated concentrations for 24 h. Cells were then trypsinized, followed by the determination of the living cell numbers by Trypan blue staining. Data are shown as means ± s.e.m., $n$ = 4 biological replicates, with $P$ values calculated by one-way ANOVA, followed by Tukey.

(**D**) Gating strategies used for quantifying the populations of dead cells. During the analysis, intact cells from each sample were selected by FSC-H and SSC-H (left, using a linear scale), followed by FSC-H and FSC-A to exclude doublets (middle, using a linear scale). The fluorescence intensities of propidium iodide (PI) and Annexin V-APC were then determined and presented as density plots (right, using a logarithmic scale). The plot was divided into four quadrants (Q1, Q2, Q3, and Q4), in which Q1 (Annexin V negative and PI positive populations) represents necroptotic cells, Q2 (Annexin V positive and PI positive populations) represents late apoptotic cells, and Q3 (Annexin V positive and PI negative populations) represents early apoptotic cells. The percentage of each type of dead cell was then calculated.

(**E** to **I**) Aldometanib does not trigger apoptosis or necroptosis in cultured HCC cells or normal liver cells. The HCC cell lines, including Huh7 (E), JHH-7 (F), Hepa1-6 (G), the primary HCC cells derived from mouse HCC tissues (H), and the normal liver cell line BNL (I), were treated with aldometanib at indicated concentrations for 2 h. Cell death was assessed using Annexin V-PI staining followed by flow cytometry. The percentages of early apoptotic, late apoptotic, and necroptotic cells are shown on the bottom of each panel as means ± s.e.m., $n$ = 3 biological replicates, with $P$ values calculated by two-way ANOVA, followed by Tukey. See also representative density plots on the top of each panel, and the gating strategy for quantifying each cell population in (D).

Experiments in this figure were performed three times.

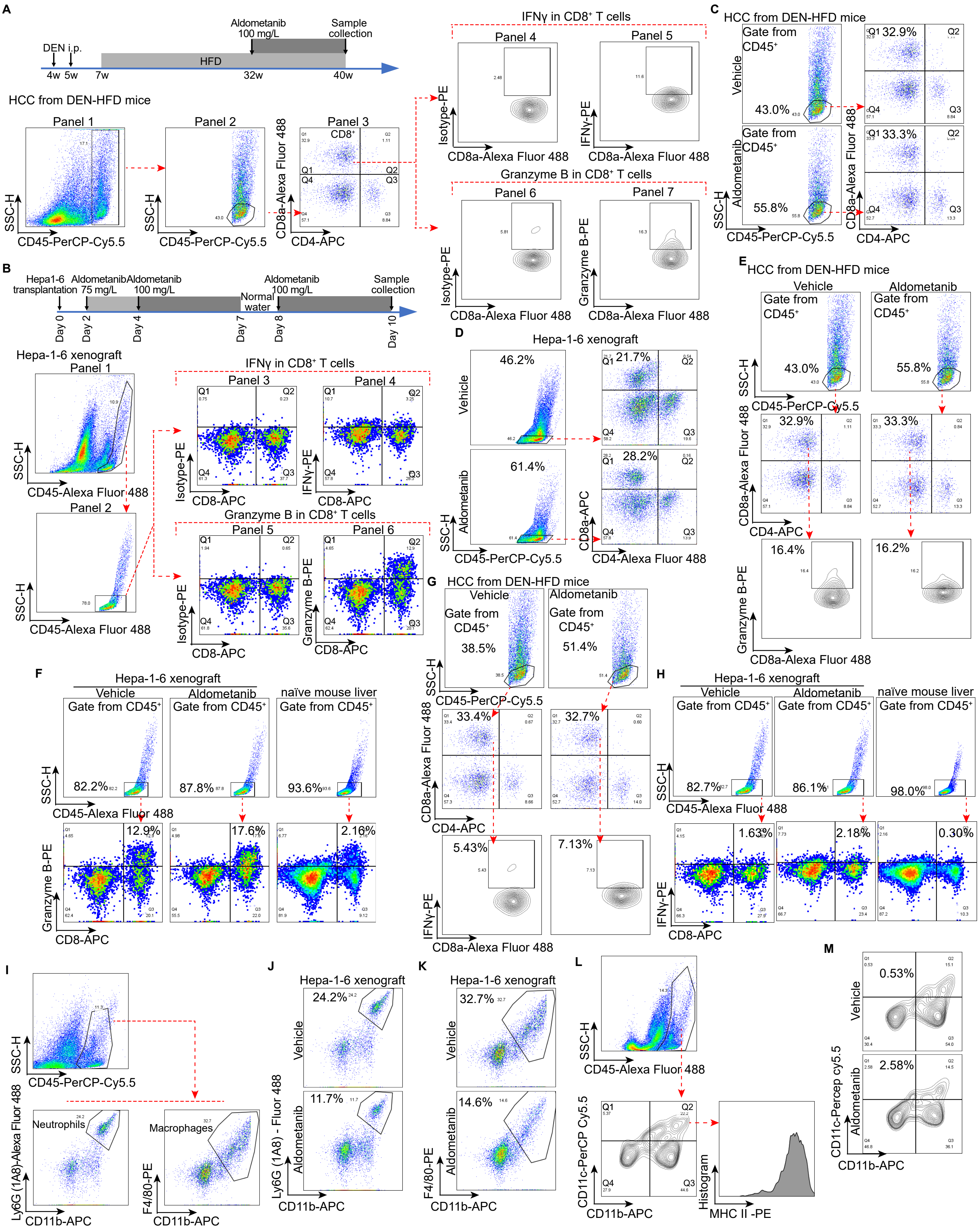
A
DEN i.p.
HFD
Aldometanib 100 mg/L
Sample collection
4w 5w 7w 32w 40w
HCC from DEN-HFD mice
Panel 1
Panel 2
Panel 3
CD8⁺
IFNγ in CD8⁺ T cells
Panel 4
Panel 5
Granzyme B in CD8⁺ T cells
Panel 6
Panel 7
SSC-H
CD45-PerCP-Cy5.5
CD8a-Alexa Fluor 488
CD4-APC
Isotype-PE
IFNγ-PE
Granzyme B-PE
C
HCC from DEN-HFD mice
Vehicle
Aldometanib
Gate from CD45⁺
43.0%
55.8%
32.9%
33.3%
B
Hepa1-6 transplantation
Aldometanib 75 mg/L
Aldometanib 100 mg/L
Normal water
Aldometanib 100 mg/L
Sample collection
Day 0 Day 2 Day 4 Day 7 Day 8 Day 10
Hepa-1-6 xenograft
CD45-Alexa Fluor 488
CD8-APC
D
Hepa-1-6 xenograft
46.2%
21.7%
61.4%
28.2%
E
HCC from DEN-HFD mice
16.4%
16.2%
F
Hepa-1-6 xenograft
naïve mouse liver
82.2%
87.8%
93.6%
12.9%
17.6%
2.16%
G
HCC from DEN-HFD mice
38.5%
51.4%
33.4%
32.7%
5.43%
7.13%
H
Hepa-1-6 xenograft
naïve mouse liver
82.7%
86.1%
98.0%
1.63%
2.18%
0.30%
I
Neutrophils
Macrophages
Ly6G (1A8)-Alexa Fluor 488
F4/80-PE
CD11b-APC
J
Hepa-1-6 xenograft
24.2%
11.7%
Ly6G (1A8) - Fluor 488
K
Hepa-1-6 xenograft
32.7%
14.6%
F4/80-PE
L
CD11c-PerCP Cy5.5
Histogram
MHC II -PE
M
0.53%
2.58%
CD11c-Percep cy5.5

**Fig. S8**

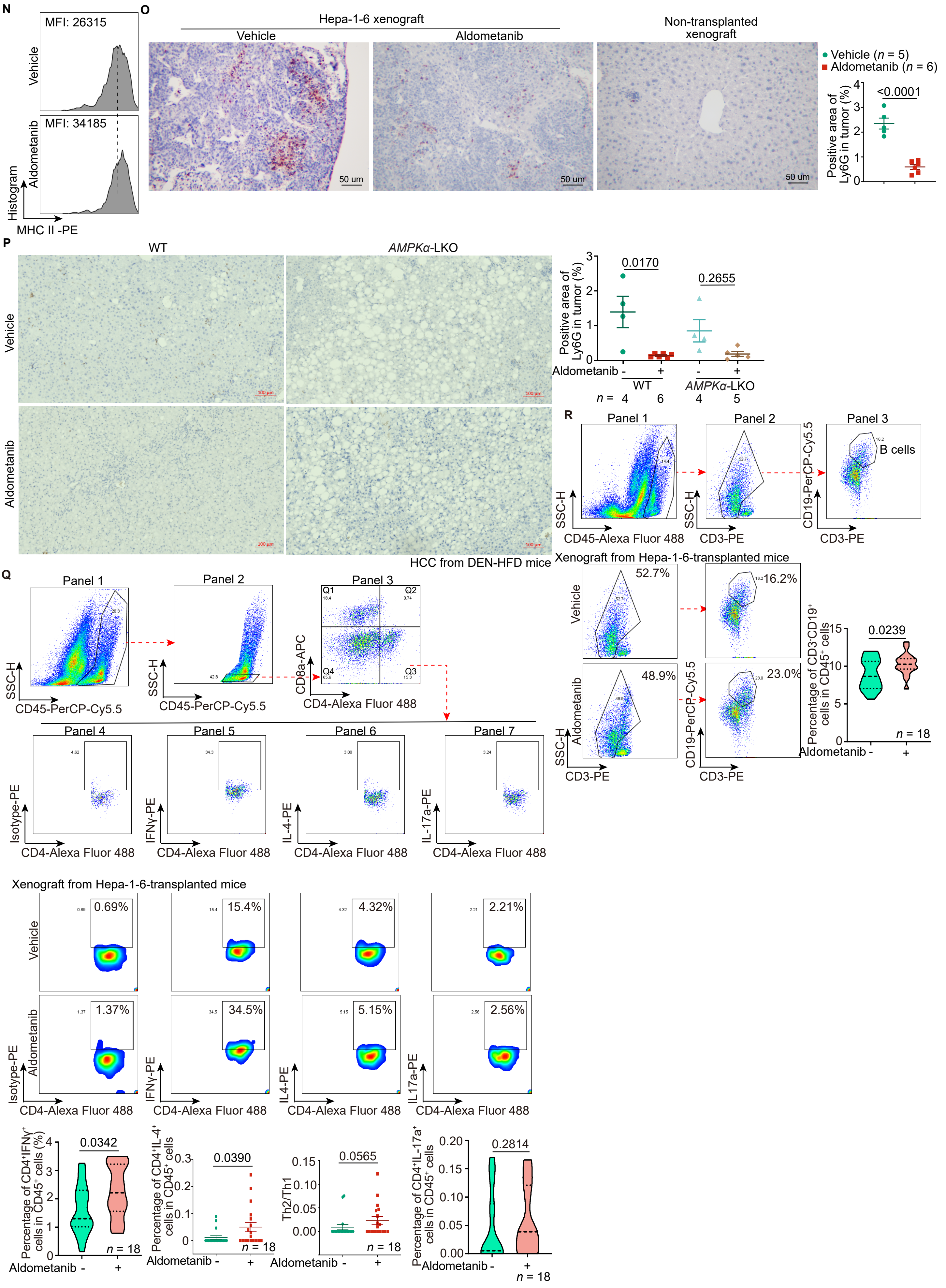

N
O
MFI: 26315
Vehicle
MFI: 34185
Aldometanib
Histogram
MHC II -PE
Hepa-1-6 xenograft
Vehicle
Aldometanib
Non-transplanted xenograft
50 um
Vehicle (n = 5)
Aldometanib (n = 6)
<0.0001
Positive area of Ly6G in tumor (%)
P
WT
AMPKα-LKO
Vehicle
Aldometanib
0.0170
0.2655
Positive area of Ly6G in tumor (%)
Aldometanib
WT
AMPKα-LKO
n =
HCC from DEN-HFD mice
R
Panel 1
Panel 2
Panel 3
B cells
SSC-H
CD45-Alexa Fluor 488
CD3-PE
CD19-PerCP-Cy5.5
Xenograft from Hepa-1-6-transplanted mice
52.7%
16.2%
48.9%
23.0%
0.0239
Percentage of CD3⁻CD19⁺ cells in CD45⁺ cells
n = 18
Q
Panel 1
Panel 2
Panel 3
CD45-PerCP-Cy5.5
CD8a-APC
CD4-Alexa Fluor 488
Q1
Q2
Q3
Q4
Panel 4
Panel 5
Panel 6
Panel 7
Isotype-PE
IFNγ-PE
IL-4-PE
IL-17a-PE
Xenograft from Hepa-1-6-transplanted mice
0.69%
15.4%
4.32%
2.21%
1.37%
34.5%
5.15%
2.56%
0.0342
0.0390
0.0565
0.2814
Percentage of CD4⁺IFNγ⁺ cells in CD45⁺ cells (%)
Percentage of CD4⁺IL-4⁺ cells in CD45⁺ cells
Th2/Th1
Percentage of CD4⁺IL-17a⁺ cells in CD45⁺ cells
n = 18


**Fig. S8 (cont.)**

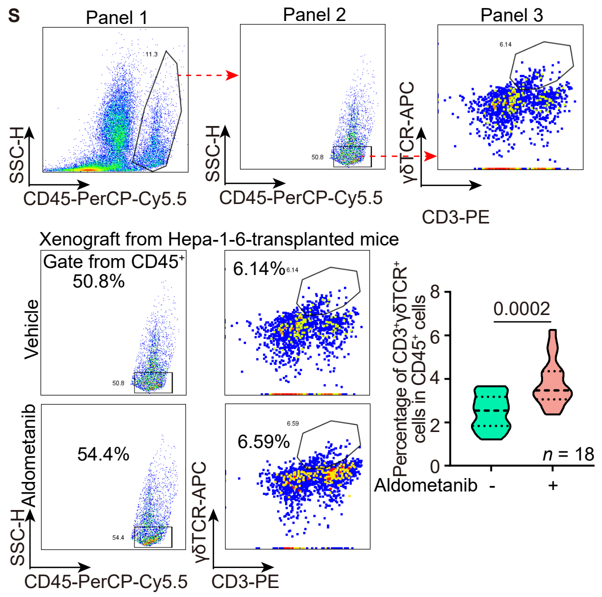


**Fig. S8 Aldometanib induces infiltration of CD8⁺ T cells into tumors.**

(**A**) Gating strategies used to quantify CD8⁺ T cells in DEN-HFD mice. HCC tissues derived from the DEN-HFD mice were digested using type I collagenase to obtain single-cell homogenates containing immune cells. The homogenates were then incubated with antibodies of CD8a-Alexa Fluor 488, CD4-APC and CD45-PerCP-Cy5.5, along with either granzyme B-PE or IFNγ-PE, or their respective isotype antibody controls that lacked specificity for the target, but matched the class and type of the antibody used to stain the target. To identify CD8⁺ T cells, the initial gating was performed using a combination of PerCP-Cy5.5 and SSC-H (panel 1), and the PerCP-Cy5.5-positive and SSC-H-low populations were selected (panel 2), which were then plotted with APC against Alexa Fluor 488 (panel 3). The cells identified as positive for Alexa Fluor 488 and negative for APC (Q1 of panel 3) represented the CD8⁺ T cells, which were then quantified. To determine the expression levels of IFNγ (panels 4 and 5) and granzyme B (panels 6 and 7) in CD8⁺ T cells, the background fluorescence of PE was first established by using CD8⁺ T cells stained with isotype control antibody (panels 4 and 6). The CD8⁺ T cells stained with granzyme B-PE or IFNγ-PE, showing PE intensities above this background (panels 5 and 7), were considered positive for PE and were then quantified. In all cytogram plots, the axes are displayed on a logarithmic scale, except for SSC-H and mean fluorescence intensity (MFI), which are shown on a linear scale; the same applies to all plots below.

(**B**) Gating strategies to quantify CD8⁺ T cells in mice transplanted with Hepa1-6 cells. Experiments were performed as in **a**, but the xenograft homogenates were not stained with CD4, and the CD45-Alexa Fluor 488 antibody was used to stain CD45. The populations that were positive for Alexa Fluor 488 with low SSC-H (panels 1 and 2) and CD8-APC (panel 3) were identified as CD8⁺ T cells. To determine the expression levels of IFNγ (panels 3 and 4) and granzyme B (panels 5 and 6) in the CD8⁺ T cells, those stained with granzyme B-PE or IFNγ-PE and showed PE intensities above the background levels (Q2 of panels 4 and 6, compared to Q2 of panels 3 and 5, which is set by the CD8⁺ T cells stained with isotype control antibody). The positive cells were then quantified.

(**C** and **D**) Aldometanib promotes infiltration of CD8⁺ T cells into HCC tissues. The HCC tissues from the DEN-HFD mice (C) or Hepa1-6-derived xenografts (D) were digested, followed by quantifying CD8⁺ T cells as in A (except that in (D), the CD45-PerCP Cy5.5, CD8a-APC and CD4-Alexa Fluor 488 antibodies were used to stain CD8⁺ T cells). Representative density plots are shown, with the percentages from gated cell populations labeled on cytograms and the same hereafter for all cytograms. The percentage of CD8⁺ T cells in the vehicle group of c, for example, is 14.1%, which is calculated by multiplying 43.0% (left panel, representing the CD45-expressing and SSC-H-low populations) by 32.9% (the Q1 of the right panel, representing the CD45-expressing and SSC-H-low populations that express CD8a but not CD4). See also statistical analysis data in Fig. 4A (C), B (D).

(**E** to **H**) Tumouricidal activity can be detected in the CD8⁺ T cells found in HCC. Experiments were performed as in Fig. 4, E to H, and the representative density plots for the populations of CD8⁺ T cells with expression of granzyme B ((E) and (F)) and IFNγ ((G) and (H)) are shown. See also the gating strategies in A ((E) and (G)) and B ((F) and (H)).

(**I** to **P**) Aldometanib eases the immune barrier in the HCC microenvironment. The quantities of neutrophils (J), macrophages (K), DCs (M), and the levels of MHC II expression in the myeloid DCs (N) were quantified by flow cytometry as described in Fig. 4I (J), J (K), K (M) and L (N). The representative density plots are shown in (J), (K), (M), and (N). The gating strategies used to quantify the neutrophils and macrophages are shown in (I), while those for DCs are shown in (L). The population that is positive for CD11b-APC and Ly6G (1A8)-Alexa Fluor 488 (I) represents neutrophils, the population that is positive for CD11b-APC and F4/80-PE (I) represents macrophages, the CD11c-PerCP-Cy5.5-positive and CD11b-negative population represents lymphoid DCs (Q1 of (L)) and the CD11c-PerCP-Cy5.5-positive and CD11b-positive population represents myeloid DCs (Q2 of (L)). To determine the expression of MHC II in myeloid DCs, the MFI (geometric mean value) of PE in the CD11b-APC- and CD11c-PerCP-Cy5.5-positive populations, as depicted in the lower right panel of (L), was further analyzed. In addition, the presence of neutrophils in both Hepa1-6 xenografts (O) and DEN-HFD-induced HCC (P) was determined by immunohistochemistry staining. Representative images are shown on the left panels, and the percentages of Ly6G-positive cells within the tumor area were calculated and are shown on the right panel (means ± s.e.m., *n* represents the number of mice, and are labeled in each panel; *P* values were calculated by two-sided Student's *t*-test (O), or two-way ANOVA, followed by Tukey (P)). The scale bars are 100 μm.

(**Q**) Aldometanib promotes infiltration of Th1 cells. The Hepa1-6-derived xenografts, collected as in (B), were digested with type I collagenase, followed by quantification of CD4⁺ T cells, including the Th1 subset (stained with CD45-PerCP-Cy5.5, CD4-Alexa Fluor 488, and IFNγ-PE antibodies), the Th2 subset (stained with CD45-PerCP-Cy5.5, CD4-Alexa Fluor 488, and IL-4-PE antibodies), and the Th17 subset (stained with CD45-PerCP-Cy5.5, CD4-Alexa Fluor 488, and IL-17a-PE antibodies), by flow cytometry. Representative density plots are shown on the middle panel, and the statistical analysis data on the lower panel (means ± s.e.m., *n* = 18 samples from 6 mice, with *P* values calculated by two-sided Student's *t*-test (left panel), or by Mann Whitney test (others). See also the gating strategies for each subset of CD4⁺ T cells on the upper panel, where the CD45-PerCP-Cy5.5-positive, SSC-H-low (panels 1 and 2), and CD4-Alexa Fluor 488-positive, CD8a-APC-negative (Q3 of panel 3) populations were identified as CD4⁺ T cells. The CD4⁺ T cells that are positive for IFNγ-PE represent the Th1 subset (panel 5), while those positive for IL-4-PE represent the Th2 subset (panel 6), and the cells positive for IL-17a-PE indicate the Th17 subset (panel 7). All these subsets exhibited higher PE intensities compared to the background PE intensities set by CD4⁺ T cells stained with isotype controls (panel 4).

(**R**) Aldometanib promotes the infiltration of B cells. Experiments were performed as in (Q), except that B cells (stained with CD3-PE, CD19-PerCP-Cy5.5, and CD45-Alexa Fluor 488 antibodies) were determined. Representative density plots are shown on the lower left panel, and the statistical analysis data on the lower right panel (means ± s.e.m., *n* = 18 samples from 6 mice, with *P* values calculated by two-sided Student's *t*-test). See also the gating strategies on the upper panel, where the Alexa Fluor 488-positive, PE-negative (panels 1 and 2), and PerCP-Cy5.5-positive (panel 3) populations were identified as B cells.

(**S**) Aldometanib promotes infiltration of γδT cells. Experiments were performed as in (Q), except that γδT cells (labeled with CD3-PE, γδTCR-APC, and CD45-PerCP-Cy5.5 antibodies) were determined. Representative density plots are shown on the lower left panel, and the statistical analysis data on the lower right panel (means ± s.e.m., *n* = 18 samples from 6 mice, with *P* values calculated by two-sided Student's *t*-test with Welch's correction). See also the gating strategies on the upper panel, where the PerCP-Cy5.5-positive (panel 1), and the PerCP-Cy5.5-positive and SSC-H-low populations were selected (panel 2), which were then plotted with PE-positive, and APC-positive (panel 3) populations were identified as γδT cells.

Experiments in this figure were performed three times.

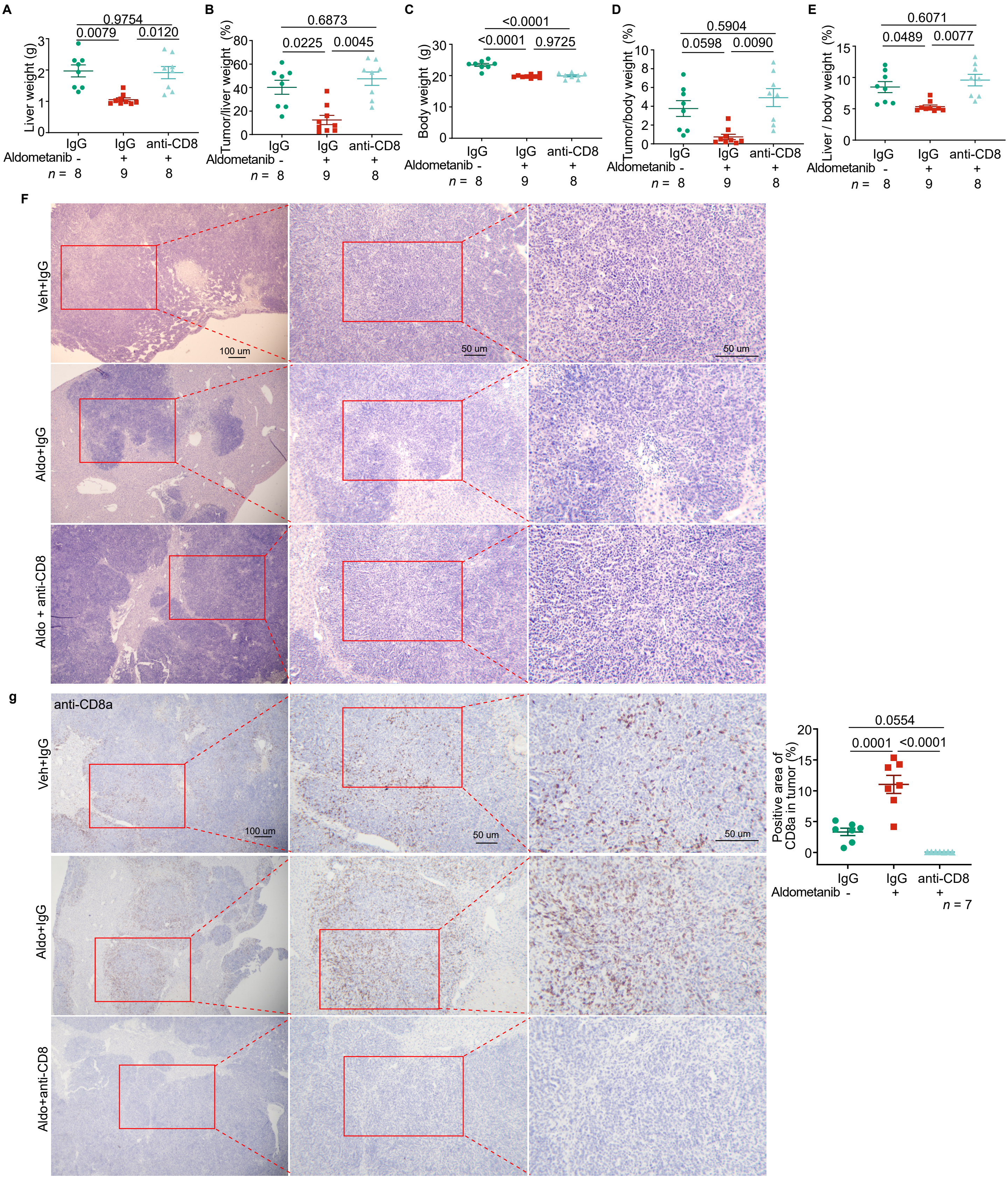


**Fig. S9 CD8⁺ T cells are required for aldometanib-induced suppression of HCC.**

(**A** to **G**) Depletion of CD8⁺ T cells diminishes the aldometanib-mediated suppression of HCC. Mice were transplanted into the left lobe with Hepa1-6 cells, treated with aldometanib, and depleted of CD8⁺ T cells, as in Fig. 4M. HCC tissue samples were collected, followed by determination of the liver weights (A), tumor:liver weight ratios (B), body weights (C), tumor:body weight ratios (D), and liver:body weight ratios (E). Data are shown as means ± s.e.m., *n* represents the number of mice, and are labeled in each panel, with *P* values calculated by two-way ANOVA, followed by Tukey), and morphology ((F); by H&E staining, and (G); by immunohistochemistry staining of CD8a, in which representative images are shown on the left, and the percentages of CD8a-positive areas within the tumor region were calculated and are shown on the right (means ± s.e.m., *n* represents the number of mice, and are indicated in each panel; *P* values were calculated by two-way ANOVA, followed by Tukey).

Experiments in this figure were performed three times.

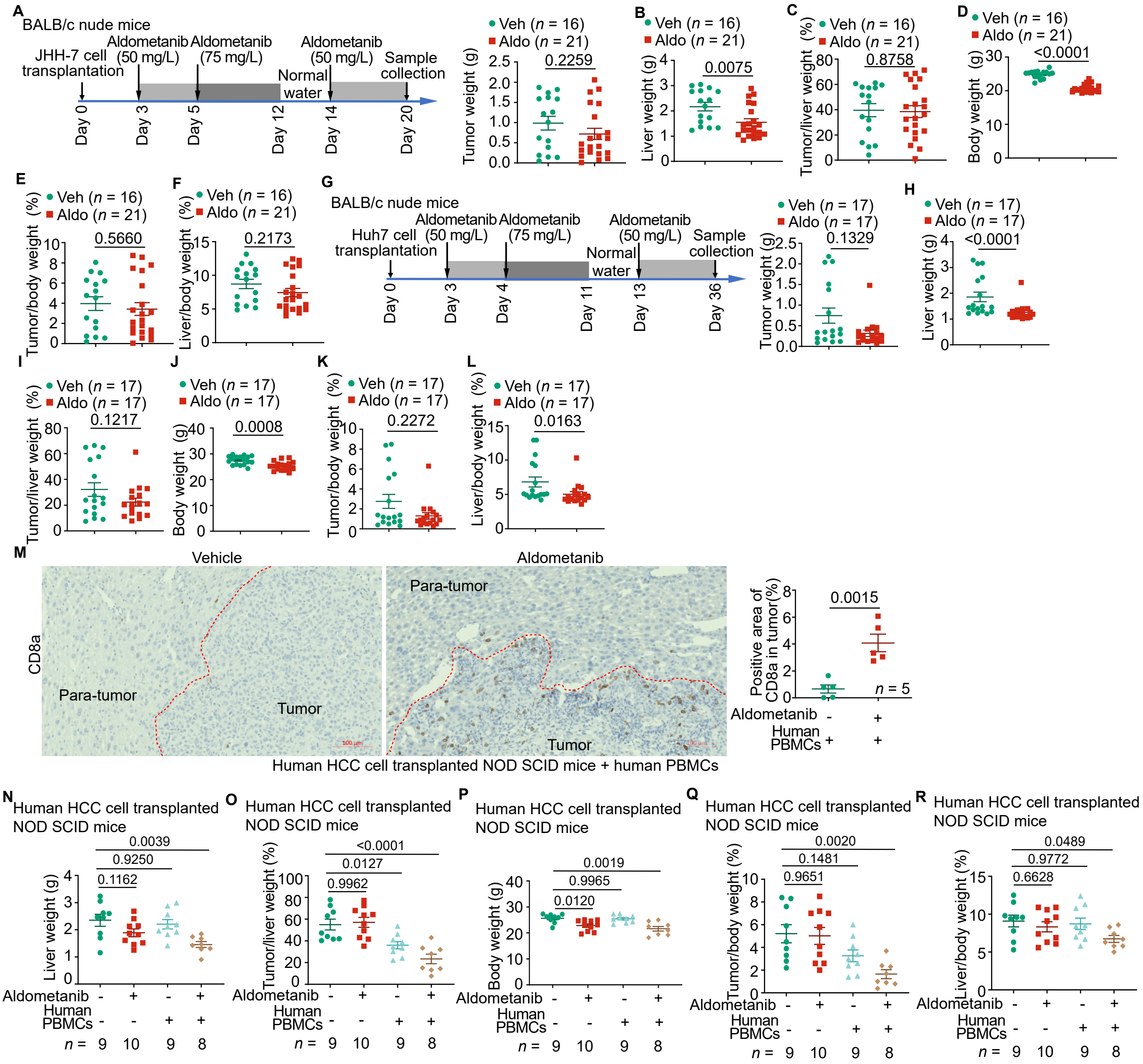


**Fig. S10 Aldometanib hardly inhibits xenografts grown in the liver of nude mice lacking CD8⁺ T cells.**

(**A** to **F**) BALB/c nude mice were transplanted with JHH-7 cells, followed by treatment of aldometanib starting from day 3 post-transplantation as described in (A). HCC tissue samples were collected on day 20, followed by determination of the tumor weights (A), liver weights (B), tumor:liver weight ratios (C), body weights (D), tumor:body weight ratios (E), and liver:body weight ratios (F). Data are shown as means ± s.e.m., *n* represents the number of mice, and are labeled in each panel, with P values calculated by two-sided Student's t-test ((A) to (E)) or by Mann-Whitney test (F).

(**G** to **L**) BALB/c nude mice were transplanted Huh7 cells, followed by treatment of aldometanib starting day 3 post-transplantation as described in (G). HCC tissue samples were collected on day 36, followed by determination of the tumor weights (G), liver weights (H), tumor:liver weight ratios (I), body weights (J), tumor:body weight ratios (K), and liver:body weight ratios (L). Data are shown as means ± s.e.m., *n* represents the number of mice, and are labeled in each panel, with *P* values calculated by two-sided Student's *t*-test (J), by two-sided Student's *t*-test with Welch's correction (I), or by Mann-Whitney test (others).

(**M** to **R**) Re-introduction of human PBMCs restores inhibitory effects of aldometanib on HCC in NOD-SCID mice. The NOD-SCID mice were intraperitoneally injected with human PBMCs, followed by transplantation with HCC cells from human HCC tissues, as in Fig. 4N. HCC tissue samples were collected, followed by determination of $CD8^+$ T cell infiltration ((M), by immunohistochemistry, staining for CD8a; representative images are shown on the left, and the percentages of CD8a-positive areas within the tumor region were calculated and are shown on the right (means ± s.e.m., *n* represents the number of mice, and are labeled in each panel; *P* values were calculated by two-sided Student's *t*-test)), liver weights (N), tumor:liver weight ratios (O), body weights (P), tumor:body weight ratios (Q), and liver:body weight ratios (R). Data are shown as means ± s.e.m., *n* represents the number of mice, and are labeled in each panel, with *P* values calculated by two-way ANOVA, followed by Tukey.

Experiments in this figure were performed three times.

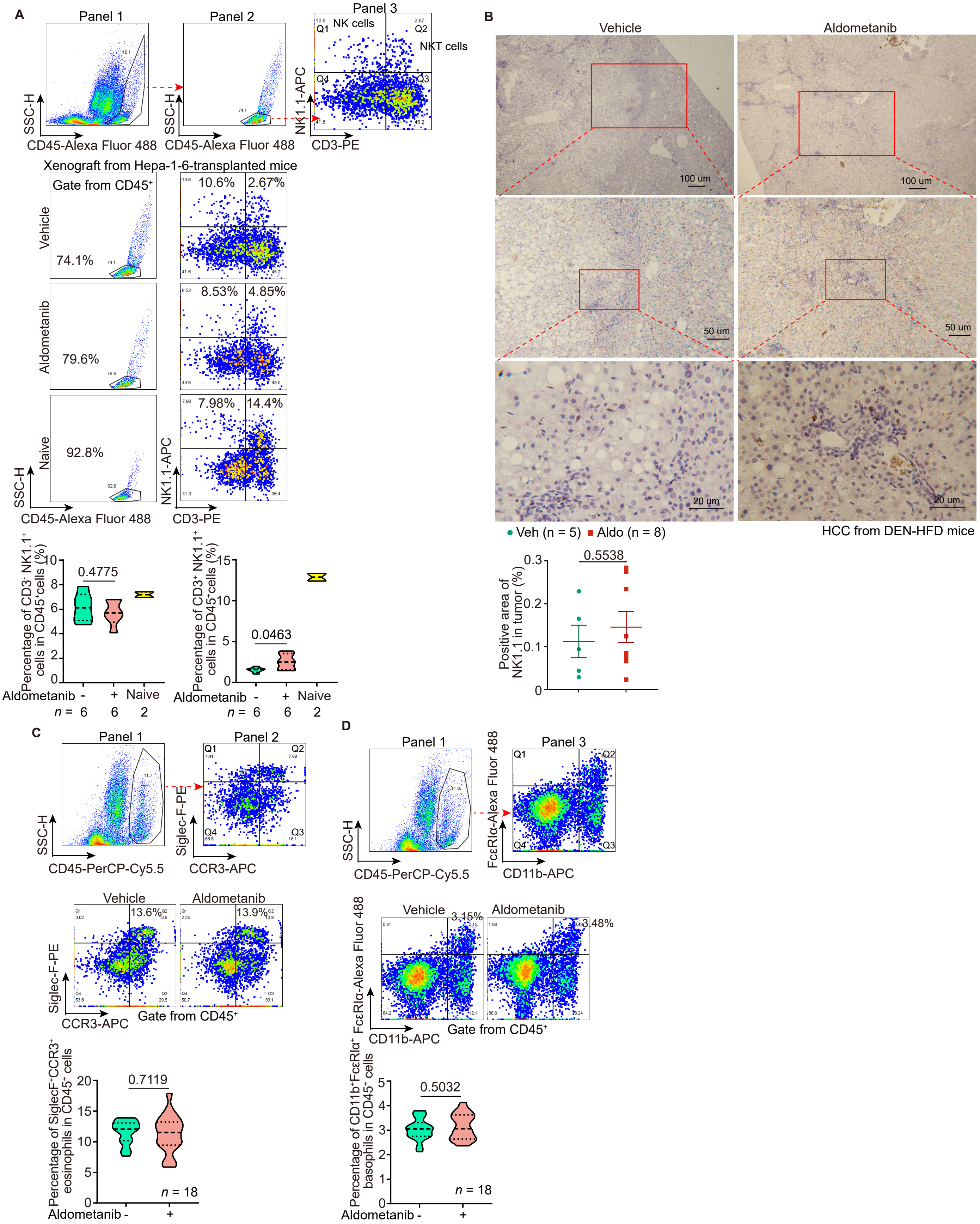


**Fig. S11 Effects of aldometanib on other tumor-infiltrating lymphocytes than CD8⁺ T cells.**

(**A** and **B**) Aldometanib promotes infiltration of NKT, but not NK cells, into HCC tissues. Experiments were performed as in fig. S8B, except that NK and NKT cells were determined using both flow cytometry ((A); with CD3-PE, NK1.1-APC, and CD45-Alexa Fluor 488 antibodies) and immunohistochemistry staining ((B); with NK1.1 antibody; representative images are shown on the upper, and the percentages of NK1.1-positive area in the tumor were calculated and are shown on the lower panel as means ± s.e.m., *n* represents the number of mice, and are labeled in each panel; and *P* values were calculated by two-sided Student's *t*-test). Representative density plots of flow cytometry are shown on the middle panel of (A), and the statistical analysis data on the lower panel of (A) (means ± s.e.m., *n* = 18 samples from 6 mice, with *P* values calculated by two-sided Student's *t*-test (the left panel), or by two-sided Student's *t*-test with Welch's correction (the right panel)). See also the gating strategies on the upper panel of (A), where the Alexa Fluor 488-positive and SSC-H-low (panels 1 and 2) cells were selected. Among these selected cells, those that were positive for PE and APC (Q2 of panel 3) were identified as NKT cells, while those negative for PE but positive for APC (Q1 of panel 3) were NK cells.

(**C** and **D**) Aldometanib does not promote infiltration of eosinophils and basophils. Experiments were performed as in fig. S8B except that eosinophils (stained with CD170-Siglec-F-PE, CCR3-APC, and CD45-PerCP-Cy5.5 antibodies) and basophils (labeled with FcεRIα-Alexa Fluor 488, CD11b-APC, and CD45-PerCP-Cy5.5 antibodies) were determined. Representative density plots are shown on the middle panel, and the statistical analysis data on the lower panel (means ± s.e.m., *n* = 18 samples from 6 mice, with *P* values calculated by two-sided Student's *t*-test). See also the gating strategies on the upper panel, where the PerCP-Cy5.5- (panel 1), APC- and PE-positive (panel 2) cells were identified as eosinophils, while the PerCP-Cy5.5- (panel 1), Alexa Fluor 488- and APC-positive (panel 3) cells were basophils.

Experiments in this figure were performed three times.

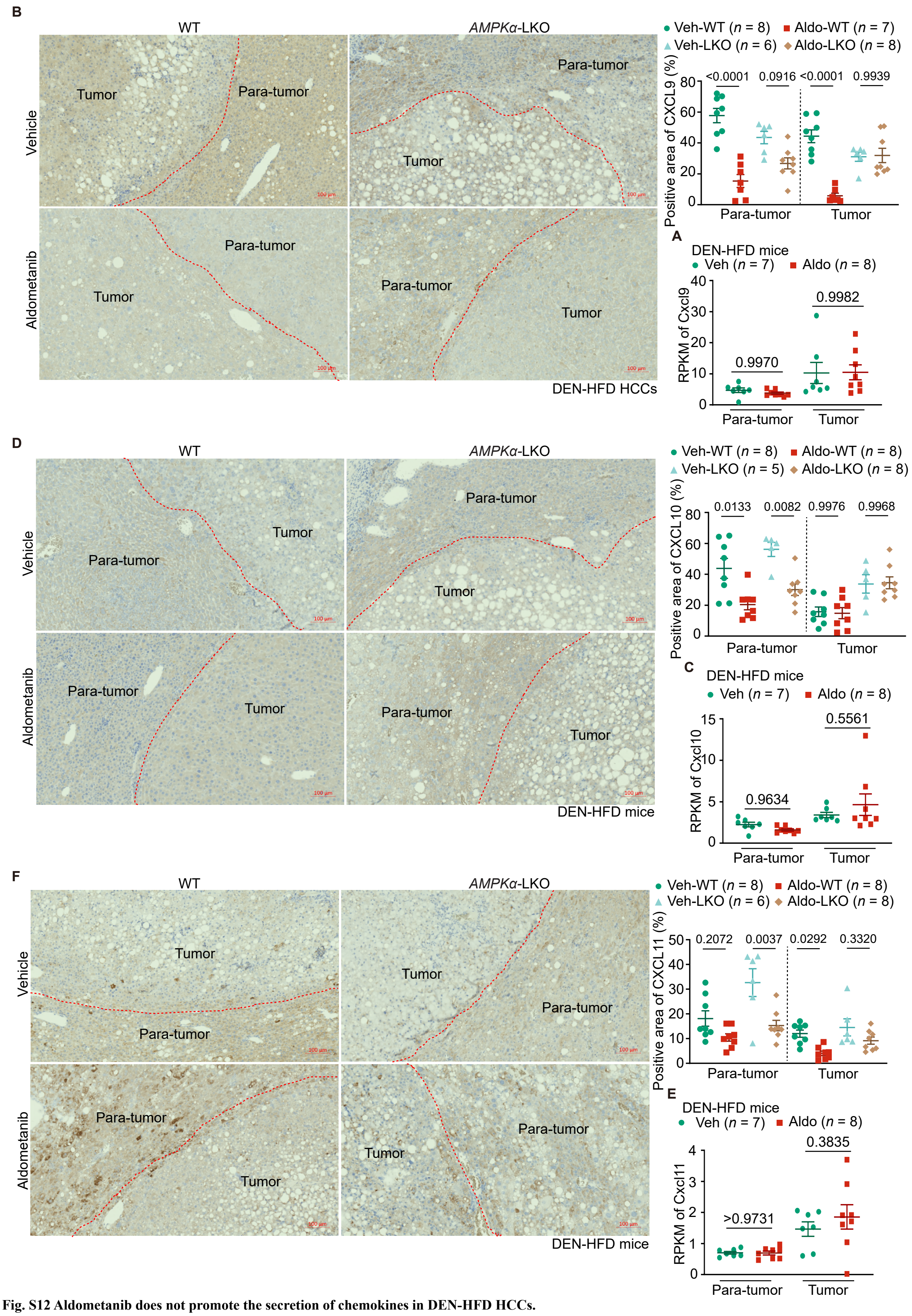


**Fig. S12 Aldometanib does not promote the secretion of chemokines in DEN-HFD HCCs.**

(**A** to **F**) Mice induced to develop HCC using DEN and HFD to week 40 of age as in fig. S8A, were treated with aldometanib as in fig. S8A. The mRNA levels ((A), (C), and (E); by RNA sequencing, shown as means ± s.e.m., *n* represents the number of mice, and are labeled in each panel, with *P* values calculated by two-way ANOVA, followed by Tukey) and protein ((B), (D), and (F); by immunohistochemistry staining; representative images are shown on the left panels, and the percentages of CXCL9/10/11-positive area within the tumor area were calculated and are shown on the right panel as means ± s.e.m., *n* represents the number of mice, and are labeled in each panel; and *P* values were calculated by two-way ANOVA, followed by Tukey) levels of CXCL9, CXCL10, and CXCL11 in both tumor and para-tumor tissues were determined. The scale bars are 100 μm.

Experiments in this figure were performed three times.

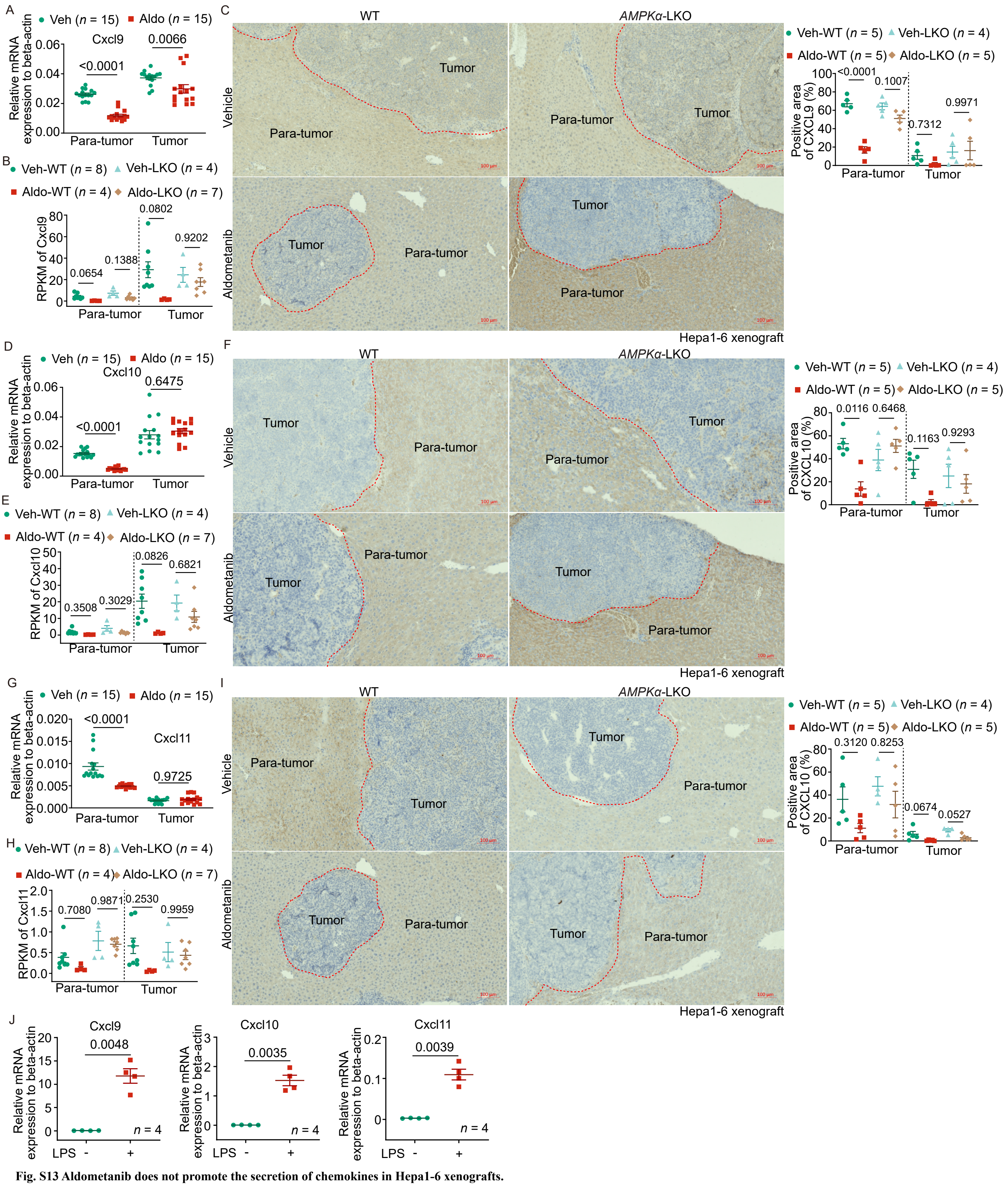


**Fig. S13 Aldometanib does not promote the secretion of chemokines in Hepa1-6 xenografts.**

(**A** to **I**) The hepa1-6-derived xenograft-bearing mice were treated with aldometanib as in fig. S8B. The chemokine (*Cxcl9*, *Cxcl10*, and *Cxcl11*) mRNA levels, determined either by RT-PCR ((A), (D), (G); shown as means ± s.e.m., *n* = 15 mice, with *P* values calculated by two-way ANOVA, followed by Tukey), or by RNA sequencing ((B), (E), (H); shown as means ± s.e.m., *n* represents the number of mice, and are labeled in each panel, with *P* values calculated by two-way ANOVA, followed by Tukey), and the chemokine protein levels determined by immunohistochemistry staining ((C), (F), (I)) in both tumor and para-tumor tissues were shown (representative images are shown on the left panels, and the percentages of CXCL9/10/11-positive area within the tumor were calculated and are shown on the right panel as means ± s.e.m., *n* represents the number of mice, and are indicated in each panel; and *P* values were calculated by two-way ANOVA, followed by Tukey). The scale bars are 100 µm.

(**J**) LPS stimulates secretion of chemokines in the liver. Wildtype C57BL/6J mice, aged 8 weeks, were intraperitoneally injected with 10 mg/kg LPS (dissolved in PBS). At 6 h after the injection, the mice were euthanized, and liver tissues were collected, followed by the determination of the mRNA levels of *Cxcl9*, *Cxcl10*, and *Cxcl11*. Data are shown as shown as means ± s.e.m., *n* = 4 mice, with *P* values calculated by two-sided Student's *t*-test with Welch's correction.

Experiments in this figure were performed three times.

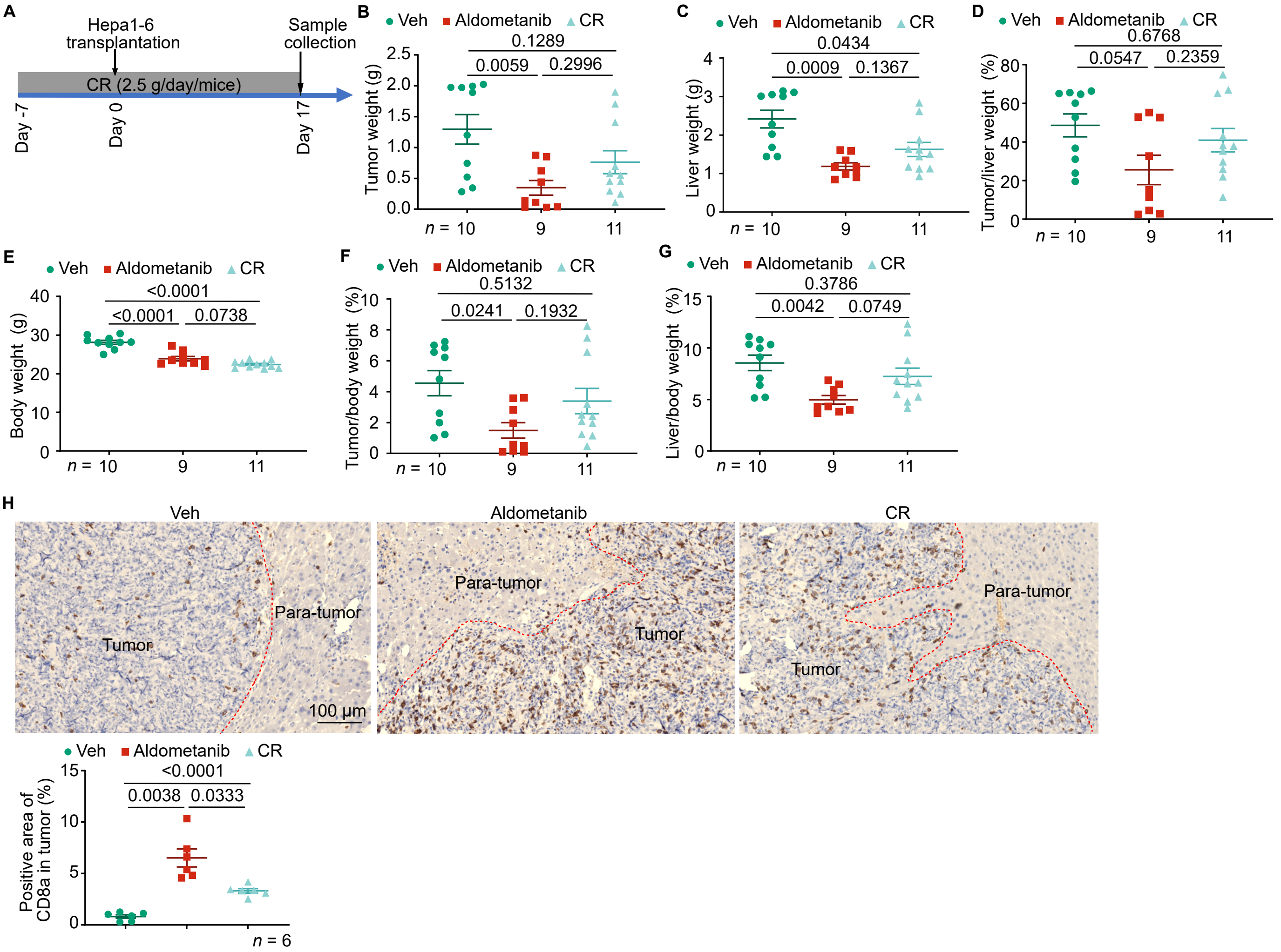


**Fig. S14 Caloric restriction suppresses growth of HCC.**

(**A** to **H**) Wildtype C57BL/6J mice, aged 8 weeks, were subjected to CR. Some 1 week after CR, Hepa1-6 cells were transplanted into the left liver lobes of mice, and the mice were further calorie-restricted for another 17 days (A). As a control, the aldometanib treatment group was set up as in Fig. 1F. The tumor weights (B), liver weights (C), tumor:liver weights (D), body weights (E), tumor:body weights (F), liver:body weights (G), and the infiltration of CD8$^{+}$ T (H), by immunohistochemistry staining for CD8a; representative images are shown on the upper panels, and the percentages of CD8a-positive areas within the tumor region were calculated and are shown on the lower panel as means ± s.e.m., *n* represents the number of mice, and are labeled in each panel; and *P* values were calculated by one-way ANOVA, followed by Tukey (B, D, E, G), or by Brown-Forsythe ANOVA test, followed by Dunnett's test (C, H). The scale bars are 100 **μm.**

Experiments in this figure were performed three times.

**Table S1 | Summary of lifespan analysis in mice[a,b]**

| Genotypes/ treatments | Mean life span (days) | | | Median life span (days) | | | | | | P-value Vs Vehicle control within each genotype (Mantel-CoX) |
|---|---|---|---|---|---|---|---|---|---|---|
| | | 95% confidence interval | | | 95% confidence interval | | | | | |
| | Estimated life span ± s.e.m. | Lower bound | Upper bound | Estimated life span ± s.e.m. | Lower bound | Upper bound | N[c] | N[d] | N[e] | |
| **Fig. 1c** | | | | | | | | | | |
| Vehicle | 563.970 ± 17.343 | 529.978 | 597.962 | 578.000 ± 27.561 | 523.980 | 632.020 | 33 | 20 | 53 | N/A |
| Aldometanib | 835.788 ± 40.509 | 756.389 | 915.186 | 805.000 ± 60.290 | 686.831 | 923.169 | 33 | 18 | 51 | N/A |

[a]Independent repeats of each lifespan experiment were performed. Data from representative experiments are shown.
[b]Lifespan data sets within each panel of this table were done in parallel and statistical analyses was done within the data set.
[c]Number of mice scored (death events).
[d]Number of mice censored.
[e]Total number of mice.
